# SugarPy facilitates the universal, discovery-driven analysis of intact glycopeptides

**DOI:** 10.1101/2020.10.21.349399

**Authors:** Stefan Schulze, Anne Oltmanns, Christian Fufezan, Julia Krägenbring, Michael Mormann, Mechthild Pohlschröder, Michael Hippler

**Author notes:** These authors contributed equally to the work (Stefan Schulze, Anne Oltmanns).

## Abstract

**Motivation:** Protein glycosylation is a complex post-translational modification with crucial cellular functions in all domains of life. Currently, large-scale glycoproteomics approaches rely on glycan database dependent algorithms and are thus unsuitable for discovery-driven analyses of glycoproteomes.

**Results:** Therefore, we devised SugarPy, a glycan database independent Python module, and validated it on the glycoproteome of human breast milk. We further demonstrated its applicability by analyzing glycoproteomes with uncommon glycans stemming from the green alga *Chlamydomonas reinhardtii* and the archaeon *Haloferax volcanii*. SugarPy also facilitated the novel characterization of glycoproteins from the red alga *Cyanidioschyzon merolae*.

**Availability:** The source code is freely available on GitHub (https://github.com/SugarPy/SugarPy), and its implementation in Python ensures support for all operating systems.

**Supplementary information:** Supplementary data are available online.

## 1 Introduction

Protein glycosylation is involved in a variety of cellular functions, ranging from protein localization and signal recognition in eukaryotes to biofilm formation and virulence in prokaryotes (Varki 2017; Eichler and Koomey 2017; Schäffer and Messner 2017). The function of a protein can be affected not only by the presence or absence of glycans but also by their type (Varki et al., eds. 2017a; Beyer et al. 2018; Di Wu et al. 2018). Therefore, the analysis of intact glycopeptides is crucial for understanding the roles of differential glycosylation in biological systems. However, this analytical task is complicated by the intricate, branched structure of this post-translational modification as well as its non-template-driven biosynthesis and compositional diversity.

Currently, commonly used engines for the analysis of intact glycopeptides rely on glycan database dependent search algorithms, e.g. Byonic™ (Bern et al. 2012), SugarQb (Stadlmann et al. 2017) and pGlyco (Liu et al. 2017b). It is immanent that such workflows can identify only glycans that are part of the respective databases. The variety of glycosylation-centric algorithms that has recently been developed (Hu et al. 2016a; Abrahams et al. 2019) also includes glycan database independent algorithms (e.g. GlycopeptideGraphMS (Choo et al. 2019), Glycoforest (Horlacher et al. 2017), glyXtoolMS (Pioch et al. 2018)). However, these tools employ graph- or network-based search strategies that still require the identification of known glycopeptides as starting points, which limits their use to studies of organisms with at least partially known glycosylation pathways. The same limitation applies to open modification search engines, which provide only mass differences, requiring additional processing to be mapped to lists of known or theoretical glycans, as well as further analysis of corresponding fragmentation spectra in order to identify glycan compositions. Nevertheless, this approach has recently been used for the characterization of bacterial glycoproteins (Ahmad Izaham, and Scott 2020). Finally, existing analysis software employing combinatorial approaches for the in silico generation of glycan libraries (e.g. GlycReSoft (Khatri et al. 2016)) are often limited in their choice of monosaccharides and/or size of the suitable search space.

Importantly, to the best of our knowledge, it remains to be shown that any of the currently available tools, including software allowing the use of customized glycan datbases (e.g. MSFragger Glyco (Polasky et al. 2020) and SugarQb (Stadlmann et al. 2017), is suitable for the analysis of large glycoproteomic datasets containing glycans with uncommon modifications (e.g. methylation) or glycans from uncharacterized glycosylation pathways. The diversity of glycosylation pathways, utilizing various monosaccharides and modifications, not only in prokaryotes but also in eukaryotes (Corfield and Berry 2015; Zhu et al. 2019; Nothaft and Szymanski 2010; Jarrell et al. 2014), thus remains largely inaccessible for high-throughput analyses. In order to close this gap, we have developed SugarPy, which is independent of glycan databases or previous information about the glycosylation pathway.

## 2 Methods

### Collection and analysis of biological samples

A detailed description for the collection and processing of all biological samples used in this study, as well as their mass spectrometric analysis, can be found in Supporting Information 1.

### Protein database search for the identification of glycopeptide sequences

Protein database search was performed by employing the search algorithms X!Tandem (version Vengeance) (Craig and Beavis 2004), MS-GF+ (version 2019.04.18) (Kim and Pevzner 2014), and MSFragger (version 20190222) (Kong et al. 2017) implemented in Ursgal (Kremer et al. 2016) by adding monosaccharides as variable modifications. Search results were statistically post-processed with Percolator (version 3.4.0) (The et al. 2016) or qvality (version 2.02) (Kall et al. 2009) and filtered by a posterior error probability ≤ 1%. Results from all engines and individual searches were merged and for the identification of *N*-glycosites, results were filtered for peptide sequences containing a modified asparagine within the consensus motif N-X-S/T. Detailed parameters used for peptide database search can be found in Supporting Information 2. It should be noted that, while this work focused on the identification of *N*-glycopeptides, peptide sequences identified that do not contain a consensus sequence could represent *O*-glycopeptides; the search engines employed are not specialized in localizing variable modifications and *O*-glycans often harbor a HexNAc at the reducing end as well.

### Design and functionality of SugarPy

SugarPy is divided into two parts, the “run” class, in which glycopeptide matching is performed, and the “results” class, in which result CSV files, elution profiles and other plots are generated. Typically, a SugarPy workflow comprises five steps:

1. Through the “run” class, Ursgal result files are parsed and peptide sequences, as well as their modifications (except monosaccharides that would be part of the glycan) and retention times (RTs), are extracted.
2. For a set of monosaccharides and maximal glycan length (both user-defined), all possible combinations of monosaccharides are calculated and the chemical compositions of the resulting theoretical glycans (taking into account glycans with the same mass) are added to the chemical compositions of the extracted set of peptides.
3. pyQms (Leufken et al. 2017) is used to build isotope envelope libraries for these theoretical glycopeptides and to match them against all MS1 spectra within the given RT windows. It should be noted that isotope envelopes consist of the theoretical *m/z* and relative intensity for all isotopic peaks. Therefore, the quality of the resulting matches is indicated by an mScore, which comprises the accuracies of the measured *m/z* and intensity.
4. For each matched molecule, a score (VL) is calculated as the length of a vector for the mScore (ranging from 0 to 1) and intensity (inorm, normalized by the maximum intensity of matched glycopeptides within the run, therefore also ranging from 0 to 1):

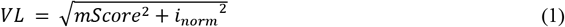 For each spectrum, all matched molecules are sorted by the glycan length (number of monosaccharides). Subsequently, starting with the longest glycan, for each glycan length, all glycan compositions are checked if they are part of any glycan composition of the previous level (longer glycan). Glycan compositions that are true subsets of larger, matched glycans (subtrees of those) are considered fragment ions and are therefore merged with the larger, final glycopeptides. It should be noted that glycan compositions can be subtrees of multiple final glycans. Furthermore, fragmentation pathways are not taken into account, however, if Y_1_-ions are matched (peptide harboring one monosaccharide), the corresponding monosaccharide is noted as the reducing end. For all final glycopeptides within one spectrum, the subtree coverage (STCov) is calculated as

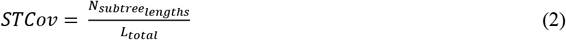

with N_subtree_lengths_ being the number of matched glycan fragment lengths (from Y_1_ to Y_n_) and L_total_ being the total length of the final glycan. Finally, the SugarPy score is calculated for each glycopeptide as the sum of vector lengths (VL, equation (1)) from all corresponding subtrees (fragment ions Y_0_ to Y_n_) multiplied by the subtree coverage (STCov, equation (2)):

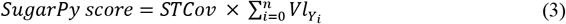 An example for the calculation of the SugarPy score is provided in Supporting Information 3.
5. SugarPy results are stored as a SugarPy “results” class in the Python pickle format. Employing functions from the results class allows to e.g. write results as CSV files, plot elution profiles (ranging from all identified glycopeptides to specific molecules, e.g. to verify that fragment ions of the same glycopeptide share the same elution profile) or plot annotated spectra.

### Application of SugarPy to biological samples

For the identification of intact *N*-glycopeptides from in-source collision-induced dissociation (IS-CID) measurements, filtered Ursgal results have been used as input files together with the corresponding mzML files. Parameters set in the Ursgal sugarpy_run_1_0_0 node for biological samples within this study can be found in Supporting Information 4. Subsequently, glycopeptide elution profiles were plotted through the sugarpy_plot_1_0_0 node, using the SugarPy result file as input.

### Filtering of SugarPy results

For automatic filtering of SugarPy results, the “extract best matches” function within SugarPy was used, accepting only the best scoring match for each spectrum and requiring a minimum of two consecutive spectra in which the glycopeptide was identified. Additionally, glycan compositions without matched Y_1_-ion were rejected and each glycopeptide was required to be found in two IS-CID runs in order to be accepted.

For manual curation, glycopeptide matches were filtered by the following criteria: (i) glycan compositions without matched Y_1_- or Y_2_-ion, were rejected; (ii) within a chromatographic elution peak, the longest glycan composition was considered; (iii) for each matched glycopeptide, one representative annotated MS1 spectrum was inspected for a reasonable Y-ion series; (iv) glycan compositions that were not in accordance with *N*-glycosylation pathways of *H. sapiens* or *C. reinhardtii* (for the respective dataset) were rejected.

For *C. merolae*, automatically filtered results were manually checked for a consistent fragmentation pattern in annotated MS1 spectra. Importantly, this did not require any prior knowledge about the *C. merolae N*-glycosylation pathway, but was solely based on the annotated glycans, as identified by SugarPy, for the respective spectra. Glycopeptides with conflicting Y-ion series or gaps of more than two monosaccharides within a Y-ion series were removed from the final results.

### Validation of glycopeptides identified by SugarPy through the analysis of NF measurements

Glycopeptides identified through the analysis of IS-CID measurements by SugarPy were checked on three levels in MS runs without IS-CID (NF measurements): (i) the presence of isotope envelopes for the corresponding glycopeptide molecules was checked in MS1 spectra using pyQms (SugarPy function “check peak presence”) and at least two isotope envelopes within a RT window of one minute surrounding the RT of IS-CID-based glycopeptide identifications was required; (ii) if MS1 isotope envelopes were found, precursors selected for HCD fragmentation within the RT windows were checked for masses that correspond to identified glycopeptides (SugarPy function “check peak presence”); (iii) for glycopeptides for which a corresponding *m/z* within the RT window was selected for HCD fragmentation, the corresponding MS2 spectra were searched for the presence of glycopeptide-specific oxonium and Y-ions. Here, the SugarPy function “check frag specs” was used, requiring a minimum of four oxonium-ions and three Y-ions, which need to include Y_1_ or Y_1_-H_2_O, and allowing a fragment ion mass tolerance of 50 ppm. The thresholds for oxonium- and Y-ions were chosen based on the number of ions that matched by chance for fragmentation spectra of peptides without glycan modification (Fig. S1).

### Data analysis and comparison with pGlyco, SugarQb, and MSFragger Glyco as well as glycan databases

pGlyco (version 2.2.2) (Liu et al. 2017b) and MSFragger Glyco (version 3.0) (Polasky et al. 2020) were integrated into Ursgal while SugarQb (Stadlmann et al. 2017) was used within Proteome Discoverer 2.1 (Thermo Scientific). Detailed parameters used to run both engines can be found in Supporting Information 5.

Human *N*-and *O*-glycan databases were downloaded from glySpace using GlycReSoft (Khatri et al. 2016) comprising 573 glycan compositions.

## Results and Discussion

SugarPy is designed for mass spectrometry (MS) data from bottom-up proteomics approaches, including enzymatic protein digestion, separation of glycopeptides by high-performance liquid chromatography (HPLC) and electrospray ionization. Furthermore, it requires separate fragmentation of peptide and glycan moiety (Fig. 1), which can be achieved through IS-CID (Hsiao and Urlaub 2010; Zhao et al. 2016). The application of higher voltages to the ion transfer region behind the source of the MS induces the fragmentation of glycosidic linkages, while the more stable peptide bonds stay intact. This glycan fragmentation results in a series of neutral losses detected in MS1 spectra, with Y-ions (nomenclature by Domon and Costello 1988) differing in the mass of one monosaccharide. The mass difference of one HexNAc can be used to dynamically select Y_0_ (peptide), Y_1_ (peptide+HexNAc) or Y_2_ (peptide+HexNAc(2)) ions for higher energy collisional dissociation (HCD) and subsequent MS2 acquisition, since Y_1_-ions are known to dominate *N*-glycopeptide fragmentation spectra (Segu and Mechref 2010). Overall, this targeting allows even low-abundant glycopeptides to be selected. Furthermore, standard protein database search engines can be used for identification of the peptide sequence from MS2 spectra by including the remaining monosaccharide(s) (HexNAc and HexNAc(2)) as variable modification. Notably, not only the peptide sequence can be determined but also the site of glycan attachment can be reliably identified. However, the glycan composition remains to be elucidated.

**Fig. 1.**
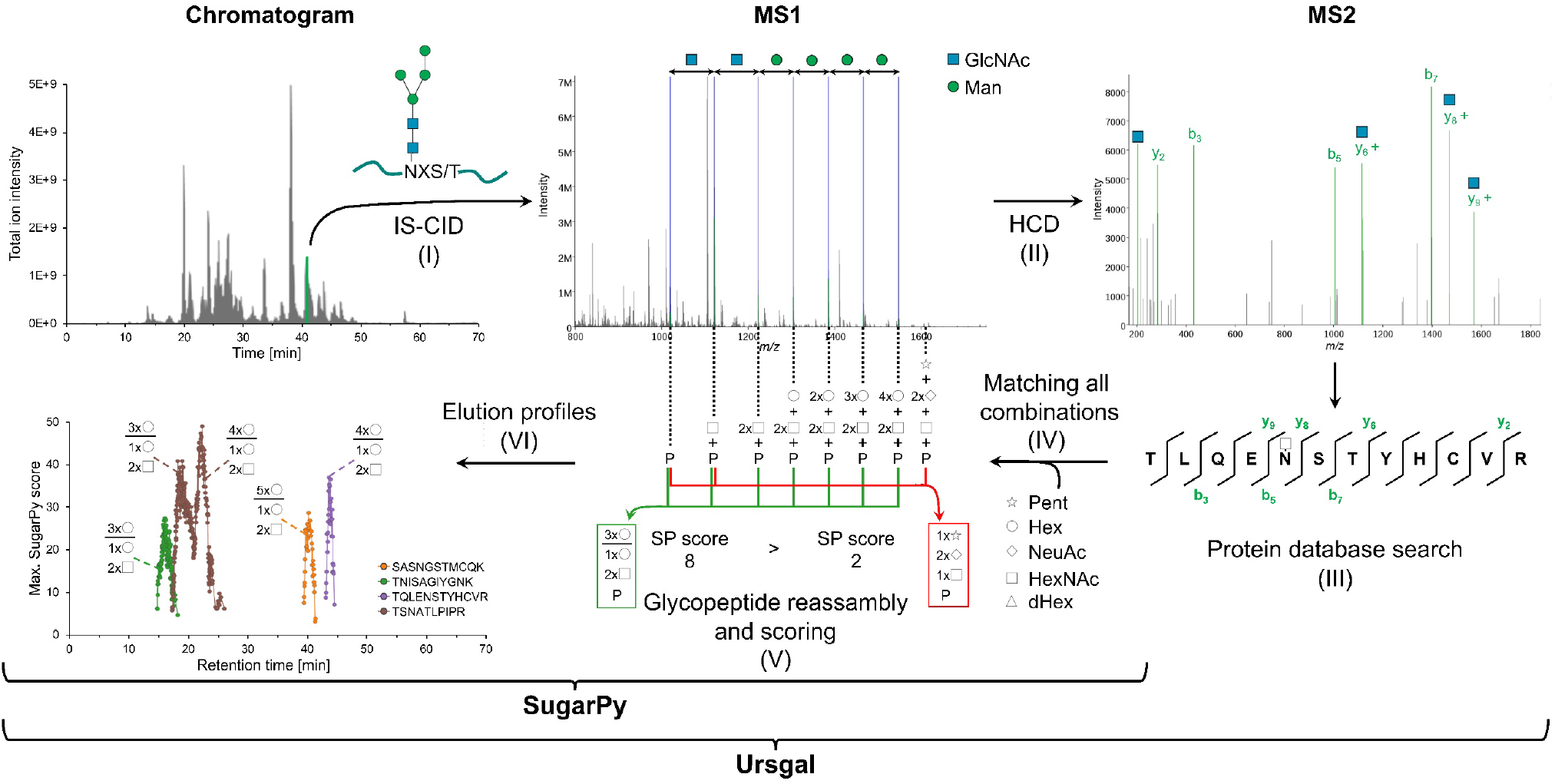
Taking advantage of IS-CID measurements, SugarPy facilitates glycan database independent analyses of intact glycopeptides. IS-CID leads to fragmentation of the glycan moiety, while the peptide backbone stays intact (I). The resulting Y-ion series detected on MS1 level can be used to dynamically target Y_0_, Y_1_ or Y_2_ ions (using mass tags) for HCD fragmentation (II). The separate fragmentation of the peptide, harboring one or two HexNAc, allows for common protein database search engines (as implemented in Ursgal) to identify the peptide sequence as well as the site of modification from MS2 spectra (III). Subsequently, SugarPy uses a set of user-defined monosaccharides to build all theoretical glycan compositions, adds them to the identified peptide sequences and matches all resulting glycopeptides to MS1 spectra within corresponding RTs (IV). Based on the detected glycopeptide fragment ions, SugarPy reassembles and scores the intact glycopeptides (V). Furthermore, corresponding elution profiles, and annotated spectra can be automatically plotted (VI). Implementation of SugarPy in Ursgal (Kremer et al. 2016), allows for high-throughput analyses of large datasets.

In order to reveal the glycan composition attached to the glycosites in the identified peptides, corresponding MS1 spectra are analyzed by SugarPy in a high-throughput discovery-driven approach (Fig. 1). For a user-defined set of monosaccharides and maximal glycan length, SugarPy computes all possible combinations of monosaccharides and adds them to peptide sequences identified from MS2 spectra, thereby creating a set of glycopeptide candidates with distinct chemical compositions. Then, the theoretical isotope envelopes of these glycopeptide candidates are matched against all MS1 spectra within the corresponding RT window of respective peptide identifications, considering entire isotope envelopes (including monoisotopic peaks) by employing pyQms (Leufken et al. 2017). From all matched candidates, SugarPy determines groups of glycan compositions that are subsets (subtrees) of a larger glycan and can therefore be regarded as corresponding glycopeptide fragment ions. The final (largest) glycopeptide is scored, taking the accuracy of isotope matching, signal intensity and number of subtrees in the respective spectrum into account. Finally, SugarPy provides elution profiles and annotated spectra (using pymzML 2.0; Kosters et al. 2018) for all scored glycopeptides allowing for a recommended manual review of the identified glycopeptides to verify e.g. the detection of expected subtrees.

As a proof of concept, we analyzed the glycoproteome of the whey fraction of human breast milk. The human *N*-glycosylation pathway is well characterized (Varki et al., eds. 2017b; Bieberich 2014) and *N*-glycoproteins are an important component of human breast milk, involved e.g. in the development of the newborn’s immune system (Zhu and Dingess 2019; Lis-Kuberka and Orczyk-Pawilowicz 2019). Recently, differential *N*-glycosylation over the course of lactation was shown with *N*-glycopeptides increasing in abundance despite a decrease in the corresponding protein level (Lu et al. 2019; Goonatilleke et al. 2019). This illustrates the need for analyses of intact glycopeptides to understand their dynamics and the molecular mechanism underlying their biological functions. After measuring human milk samples employing IS-CID, a protein database search on MS2 level resulted in the identification of 19 *N*-glycosites harboring one or two HexNAc. Using SugarPy for the automatized matching of glycopeptides in MS1 spectra yielded 211 potential glycopeptides (Fig. 2A). These initial hits were filtered based on two approaches:(i) automatic filtering, which accepted only top scoring glycopeptides identified in at least two consecutive spectra and required identification in two technical replicates, and (ii) manual filtering assessing the SugarPy score, agreement with known glycosylation pathways, and the Y-ion series in annotated MS1 spectra. In total, this IS-CID/SugarPy approach led to the identification of 20 and 36 *N*-glycopeptides that passed automatic and manual filtering, respectively.

**Fig. 2.**
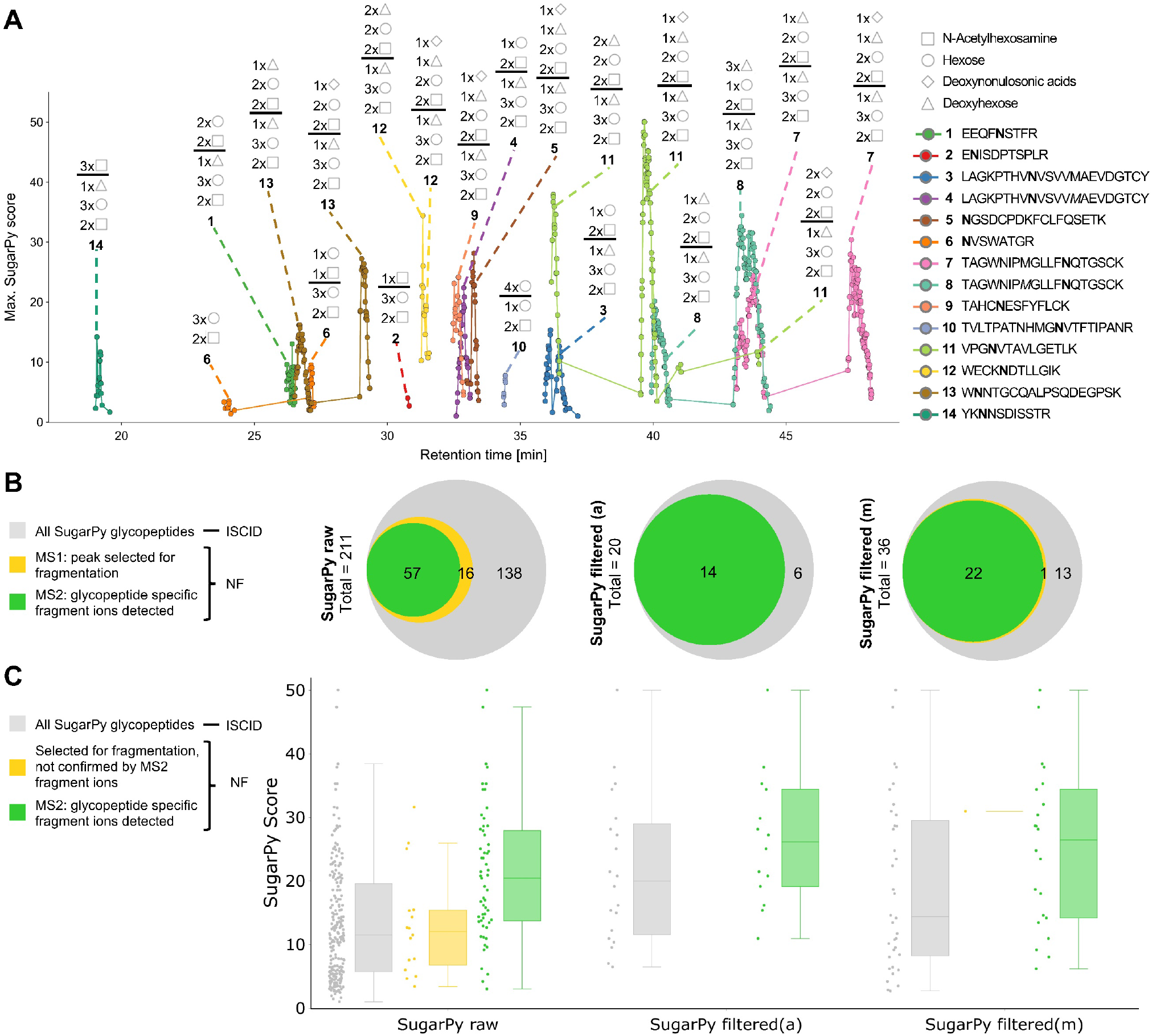
Analyses of intact *N*-glycopeptides after tryptic digest of human milk demonstrate reliable *N*-glycopeptide identifications by SugarPy. (A) A representative elution profile of *N*-glycopeptides identified by SugarPy from IS-CID measurements is shown. Traces represent the maximum SugarPy score of all *N*-glycopeptides for each MS1 spectrum for the corresponding glycopeptide sequence, but glycans are annotated only if they were accepted after manual filtering. While SugarPy reports glycan compositions, the depicted symbol representations (following the Symbol Nomenclature for Glycans (Neelamegham et al. 2019) here as well as in all other figures) separate monosaccharides that can be attributed to the glycan core (below horizontal line, based on MS1 fragmentation pattern) from the remaining monosaccharides. *N*-glycosites identified by protein database search are shown in bold letters, oxidation of methionine is indicated by italic letters. Peptide sequences correspond to the following proteins (UniProt IDs): P01859 (1), P01591 (2), P01876 (3/4), P02788 (5, 7/8), P08571 (6), P22897 (9, 12), P01024 (10), P01833 (11, 13), P01871 (14). (B), Initial raw results (left) as well as automatically (middle) and manually (right) filtered SugarPy results based on IS-CID measurements (grey) were evaluated through NF measurements. The presence of isotope envelopes on MS1 level corresponding to identified *N*-glycopeptides, their selection for HCD fragmentation (orange) and the detection of glycopeptide-specific fragment ions on MS2 level (green) is shown as area-proportional Venn diagrams. It should be noted that all *N*-glycopeptides for which isotope envelopes were detected on MS1 level were also fragmented. The corresponding areas are therefore not visible. (C), The SugarPy score distribution of all identified, unfiltered *N*-glycopeptides (left), automatically (middle), and manually (right) filtered results is represented as boxplots. For each category, the set of all corresponding *N*-glycopeptides (grey), as well as *N*-glycopeptides that were selected for HCD fragmentation and either lacked (orange) or showed (green) the presence of corresponding oxonium and Y-ions on MS2 level, is shown.

Since IS-CID fragments glycopeptides before the MS1 scan, the *m/z* of the precursor, i.e. the intact glycopeptide, is unknown and therefore, incomplete glycopeptides might be assigned. Hence, in order to verify glycopeptide identifications from the IS-CID/SugarPy approach, additional non-IS-fragmenting measurements (NF) were performed on the same samples. In such regular HPLC-MS/MS runs, glycopeptides can be detected on two levels: (i) as elution peak on MS1 level within the expected RT window and (ii) by characteristic oxonium- and Y-ions of HCD-fragmented glycopeptides on MS2 level. Indeed, MS1 elution peaks and glycopeptide-specific fragment ions were detected for 14 automatically filtered and 22 manually filtered glycopeptides identified by SugarPy (Fig. 2B). The raw SugarPy results contained 57 out of 211 glycopeptides that could be validated on MS2 level in NF runs, indicating that some glycopeptides were identified correctly by SugarPy but were removed by the filtering. However, 16 glycopeptides from the initial SugarPy hits were selected for HCD fragmentation, but the threshold of four oxonium- and three Y-ions (including Y_1_ or Y_1_-H_2_O) was not passed in the corresponding MS2 spectra indicating a potential for false positive identifications. Notably, a positive correlation between the SugarPy score and the confirmation of identified glycopeptides by NF runs was observed, i.e. for glycopeptides with higher SugarPy scores glycan-specific fragment ions were more likely to be detected in NF runs (Fig. 2C). This suggests that the SugarPy score could be used as a measure of reliability for the identified glycopeptides. Importantly, since the automatic as well as the manual filtering removed all but one glycopeptide that, although it was fragmented, was lacking glycan-specific ions, both filtering approaches were regarded as satisfactory to provide final results. However, it is worth mentioning that automatic filtering is sufficient for the reliable identification of glycopeptides from SugarPy results.

It should be noted that, while different approaches have been discussed (Hu et al. 2016b; Liu et al. 2017a; Zeng et al. 2016; Park et al. 2016; Zhu et al. 2014), a universal and accurate calculation of false discovery rates for intact glycopeptides is currently not available and therefore beyond the scope of this work. Furthermore, glycopeptides often show low intensities due to their low abundance and poor ionization properties. Therefore, detection of these glycopeptides is vulnerable to ion suppression and interference, which likely explains why 30% and 36% of the automatically and manually filtered glycopeptides, respectively, were neither detected on MS1 nor on MS2 level in NF runs. However, this does not indicate false positive matching, especially since their SugarPy score distribution is similar to the other categories. Moreover, all *N*-glycosites identified here as well as several of the identified *N*-glycopeptides have been reported previously (Table S1). Another indication for the efficient removal of false positives by the applied filtering is provided by the identified glycan compositions. Since all theoretical combinations of monosaccharides have been included in the SugarPy search, glycan compositions could be matched that are not in agreement with the well-studied human glycosylation pathways. Importantly, while 24% of glycan compositions included in the SugarPy raw results were not found in human glycan databases, all of these potential false positives were removed by the applied automatic filtering.

SugarPy results stemming from IS-CID measurements were also compared to glycopeptide identifications from the established glycopeptide search engines pGlyco (Liu et al. 2017b) and SugarQb (Stadlmann et al. 2017) as well as the recently introduced MSFragger Glyco (Polasky et al. 2020). These tools were run using existing human glycan databases and the IS-CID-independent (NF) analysis of human milk glycoproteins. In total, 24 out of 45 glycopeptides identified by SugarPy (combining both filtering approaches) were also identified by one of the three glycan database dependent search engines (Fig. S2). This overlap is another indication of the validity of the filtered SugarPy results. In addition, 28 glycopeptides (out of 166) that were excluded by the filtering, were identified by pGlyco, SugarQb or MSFragger Glyco as well, confirming again that the SugarPy raw results contain additional true positives. However, preventing the loss of these additional identifications, e.g. by more extensive manual validation of fragmentation spectra or additional replicate measurements, is beyond the scope of this manuscript. Furthermore, the large percentage of unique identifications (on the level of glycopeptides as well as glycosites) from each engine illustrates that the different approaches can complement each other. Therefore, similar to established methods for protein database searches (Jones et al. 2009; Kremer et al. 2016; Barsnes and Vaudel 2018), the combination of multiple algorithms and approaches will allow for the most comprehensive analyses. In fact, despite the relatively small size of this proof of concept dataset, the SugarPy/IS-CID approach resulted in the identification of several human milk *N*-glycopeptides that have not been described previously (Table S1).

In contrast to mammalians, some unicellular eukaryotes like *Chlamydomonas reinhardtii* synthesize *N*-glycans that, unlike human or plant glycans, contain methylated mannose (MeMan) as well as xylose (Xyl) and fucose (Fuc) residues (Mathieu-Rivet et al. 2013). Therefore, we employed SugarPy for the *N*-glycoproteomic analysis of *C. reinhardtii*, following the same approach as for the human dataset. In total, 67 *N*-glycosites were identified from IS-CID data employing Ursgal (Fig. S3). After automatic and manual filtering, 67 and 106 *N*-glycopeptides were identified by SugarPy, out of which 46 and 72 were verified, respectively, by glycopeptide-specific fragment ions in MS2 spectra of NF measurements (Fig. S4). It should be noted that chemically isomeric combinations of monosaccharides (MeMan + Xyl = Man + Fuc), led to a higher number of likely false positives in the SugarPy raw results. Therefore, we had a closer look at the annotated MS1 spectra (after IS-CID) of those seven glycopeptides that passed the automatic filtering but for which no glycopeptide-specific fragment ions in NF runs could be identified. For three out of seven glycopeptides, inconsistent or incomplete Y-ion series were revealed (Fig. S5), indicating that these are indeed false positives. While such a manual check is thus recommended (and facilitated by the automatic annotation of spectra), these instances account for less than 5% of the total *N*-glycopeptide identifications after filtering. Therefore, the applied filtering still resulted in the reliable identification of intact *C. reinhardtii N*-glycopeptides harboring uncommon modifications. The respective glycan compositions are largely in agreement with previous independent analyses using MALDI-TOF, Western blots and glycosyltransferase mutants (Mathieu-Rivet et al. 2013; Oltmanns et al. 2020b). Nevertheless, isomeric monosaccharides or monosaccharide combinations should be taken into account when interpreting results, especially since microalgae have been shown to exhibit methylation of Pent as well (MePent and dHex are isomers) (Mocsai et al. 2020).

Furthermore, comparing SugarPy results to analyses with pGlyco, SugarQb, and MSFragger Glyco highlights the limitations of glycan database dependent search algorithms. As pGlyco uses a fixed *N*-glycan database, only a small subset of *N*-glycopeptides, consisting of oligomannosidic *N*-glycans, is identified when evaluating NF measurements of *C. reinhardtii* (Fig. S6). In contrast, SugarQb and MSFragger allowed the use of a custom glycan database with the same search space as SugarPy. However, the differences are striking. For 39% and 58% of glycopeptides identified by SugarQb and MSFragger Glyco, respectively, improbable glycan compositions were matched, i.e. glycans containing ≠2 HexNAc, conflicting with the described *N*-glycosylation pathway) (Mathieu-Rivet et al. 2013). This was the case for only five glycopeptides identified by SugarPy, some of which showed glycopeptide-specific fragment ions in NF-runs and could potentially be explained by *O*-glycosylation, a pathway that has only been scarcely studied in *C. reinhardtii* so far (Mathieu-Rivet et al. 2017).

These results show that neither pGlyco nor SugarQb or MSFragger are well suited for the analysis of intact *N*-glycopeptides from *C. reinhardtii*.In contrast, the IS-CID/SugarPy approach resulted in compelling identifications. Equally, SugarPy was successfully employed in previous glycoproteomics analyses of *C. reinhardtii N*-glycosylation mutants (Schulze et al. 2018; Oltmanns et al. 2020b) as well as the characterization of novel *N*-glycopeptides from *Botryococcus braunii* (Schulze et al. 2017). In these studies, the plausibility of identified *N*-glycan compositions was confirmed through Western blots with glycan-specific antibodies as well as genomic and reverse genetics analyses of the *N*-glycosylation pathway.

While protein glycosylation is a common feature of all domains of life, a common *N*-glycan core is missing for most prokaryotes. Instead, a diverse array of monosaccharides is used in a multitude of glycosylation pathways (Schäffer and Messner 2017; Jarrell et al. 2014) rendering prokaryotic glycans largely inaccessible to glycan database dependent search engines. However, when employing SugarPy for the evaluation of IS-CID measurements of the archaeon *Haloferax volcanii, N*-glycan compositions in accordance with literature were obtained after automatic filtering (Fig. S7) (Esquivel et al. 2016). This holds true even though SugarPy was parameterized with a set of 12 different monosaccharides (>70,000 theoretical combinations). Furthermore, in measurements of a mutant strain lacking the oligosaccharyltransferase AglB, as expected, no glycopeptides were identified (Fig. S7B). This illustrates again the reliability and broad applicability of the IS-CID/SugarPy approach.

Finally, SugarPy facilitated the analysis of *N*-glycoproteins from the acidophilic, thermophilic red alga *Cyanidioschyzon merolae*. Bioinformatic analyses of the *C. merolae* genome indicated that an *N*-glycosylation pathway is present (Oltmanns et al. 2020a). However, the complexity of this *N*-glycosylation pathway and consequently the *N*-glycan compositions of *C. merolae* were unknown. Therefore, we defined a broad set of eukaryotic monosaccharides in SugarPy. Results were automatically filtered, thereby preventing any need for prior knowledge about the glycan composition. In addition, annotated MS1 spectra of filtered results were manually checked for continuous Y-ion series, providing additional confidence as indicated by the analysis of *C. reinhardtii* results (see above). In total, 15 intact *N*-glycopeptides were identified, harboring mainly oligomannosidic and partly methylated *N*-glycans with four to ten hexoses. Interestingly and contrasting other microalgae, *N*-glycopeptides from *C. merolae* identified by SugarPy were not found to be decorated with Fuc or Xyl (Fig. S8A,B). This finding was confirmed by subjecting samples to immunoblotting using Fuc and Xyl specific antibodies resulting in no detectable signal for either antibody (Fig. S8C).

## 4 Conclusion

In conclusion, with SugarPy we have successfully developed a glycan database independent search algorithm that can determine glycan compositions using a de novo approach. Together with common bottom-up proteomics tools used to determine the peptide sequence, the IS-CID/SugarPy approach facilitates the universal analysis of intact glycopeptides. We tested our algorithm on intact glycopeptides synthesized by the known *N*-glycosylation pathways of human (milk whey fraction) as well as *C. reinhardtii* (culture supernatant). Both confirmed the reliability of SugarPy for the identification of *N*-glycopeptides, as filtered SugarPy results were validated by glycopeptide-specific fragment ions in MS runs without IS-CID. While manual filtering of SugarPy raw results increases the number of identifications, automatic filtering is sufficient to yield reliable identifications and can be combined with a manual check of annotated spectra without requiring prior knowledge about the *N*-glycosylation pathway. The broad applicability of this discovery-driven approach facilitated by SugarPy is emphasized by the analysis of *N*-glycopeptides of the haloarchaeon *H. volcanii*, and the characterization of novel *N*-glycopeptides of the red algae *C. merolae*. Although these analyses were focused on *N*-glycopeptides, the generalized design of SugarPy allowed for the identification of *O*-glycopeptides in a human data set as well (Fig. S9). Finally, while SugarPy is currently designed for IS-CID data, other fragmentation techniques like EThcD (Yu et al. 2017), that separate glycan from peptide fragmentation, could be analyzed with SugarPy. Even its application to combined fragmentation spectra is plausible, given the fragmentation method provides a high Y-ion series coverage. SugarPy is publicly available (https://github.com/SugarPy/SugarPy) and integrated into the Python framework Ursgal (Kremer et al. 2016), allowing for straightforward integration into analysis workflows.

### Code and data availability

The source code for SugarPy is available through GitHub (https://github.com/SugarPy/SugarPy). Example scripts to replicate the workflows presented here, as well as a script to plot annotated spectra for any identified glycopeptide is provided.

All MS raw data that have been generated or used within this study have been deposited to the ProteomeXchange Consortium (Deutsch et al. 2017) via the PRIDE partner repository (Perez-Riverol et al. 2019) with the dataset identifier PXD017345. This includes Ursgal result files, SugarPy result files as well as results from SugarQb and pGlyco.

## Acknowledgements

We would like to thank Martin Scholz for support in setting up IS-CID methods on the MS, helpful discussions as well as for suggesting SugarPy’s name. Furthermore, we thank Prof. Dr. Antje von Schaewen for providing *A. thaliana* mutant lines. Funding from the German Research Foundation for S.S. (DFG grant 398625447) and M.H. (HI 739/12-1) is greatly acknowledged. M.H. acknowledges further funding by the Sino-German Center, Beijing, China (project GZ990). M.P. was supported by the National Science Foundation grant 1817518.

## Author contributions

SS, CF and MH devised the concept for this work. SS developed SugarPy together with CF. AO and JK performed sample preparation and MS measurements, SS and AO wrote the manuscript supported by CF, MP and MH. MP, MH and MM supervised the work.

