## Supporting information for "SugarPy facilitates the universal, discovery-driven analysis of intact glycopeptides"

[1]University of Pennsylvania, Department of Biology, Leidy Laboratories, Philadelphia, USA, [2]University of Muenster, Institute of Plant Biology and Biotechnology, Muenster, Germany, [3]Heidelberg University, Institute of Pharmacy and Molecular Biotechnology, Heidelberg, Germany, [4]University of Muenster, Institute for Hygiene, Muenster, Germany, [5]Institute of Plant Science and Resources, Okayama University, Kurashiki, Japan

[†]These authors contributed equally to the work

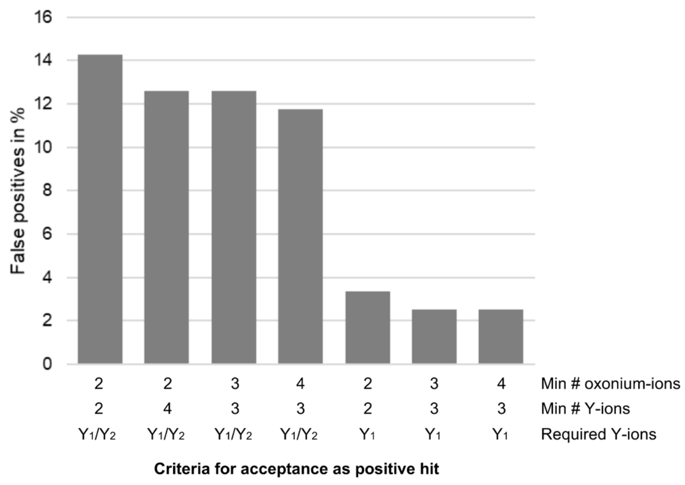


**Fig. S1. A minimum of four oxonium- and three Y-ions (including $Y_1$) for the matching of glycopeptide-specific fragment ions in MS2 spectra shows the lowest rate of potential false positives.** The rate of potential false positives has been determined in an non-IS-fragmenting MS run of tryptic human milk (glyco-)peptides. Peptide sequences without glycan modification were determined from MS2 spectra by protein database search and filtering results for PEP <= 1% (see Supporting Information 2). For each identified peptide, respective MS2 spectra were searched for glycopeptide-specific fragment ions corresponding to the peptide harboring the randomly chosen glycan Hex(5)HexNAc(3)NeuAc(1)dHex(1). Matched ions were considered false positives since the spectra were determined to originate from non-glycosylated peptides. The percentage of peptides (from a total of 119), for which false positive glycopeptide-specific ions were detected, is given for different thresholds of oxonium- and Y-ions (including required $Y_1$- or $Y_2$-ions). It should be noted that these rates are most likely higher than the respective ones for the validation of glycopeptides identified by SugarPy, because the latter includes elution within the same RT window as well as matching MS1 precursor isotope envelopes as additional requirements.

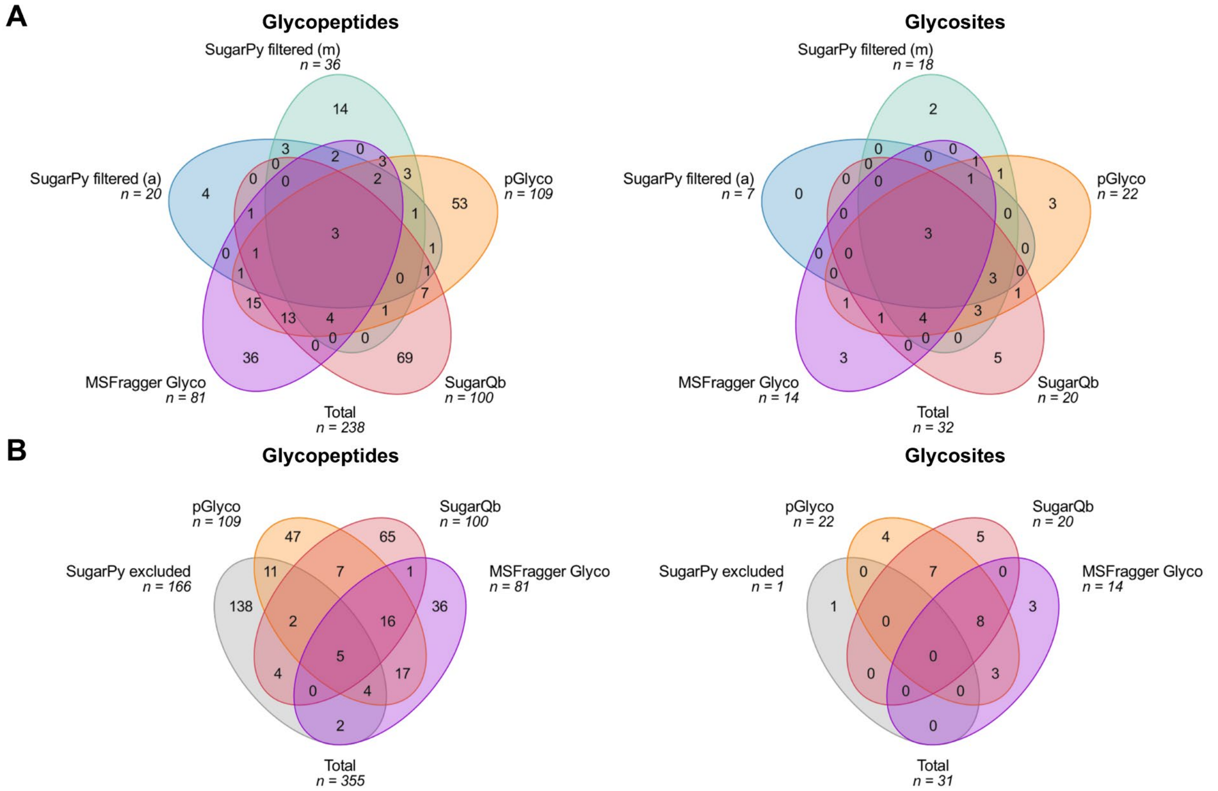


**Fig. S2. Comparison of *N*-glycopeptide identifications from SugarPy, pGlyco, SugarQb and MSFragger Glyco shows unique identifications for each search engine.** (**A**), Venn diagrams for *N*-glycopeptides (defined by the combination of peptide sequence, modification and glycan composition) identified by the IS-CID/SugarPy approach with automatic (blue) and manual (green) filtering or by pGlyco (orange), SugarQb (red) and MSFragger Glyco (purple) from NF measurements, illustrate a high degree of unique identifications for each engine. (**B**), The comparison of *N*-glycopeptides that were rejected by automatic and manual filtering (grey) with pGlyco, SugarQb and MSFragger Glyco results indicates that some probable *N*-glycopeptide identifications were removed from SugarPy results by filtering. (**C**) and (**D**), The Venn diagrams of *N*-glycosites for the same categories as in A and B, respectively, illustrates that the IS-CID/SugarPy approach can identify unique *N*-glycosites, while a lower number of *N*-glycosite identifications through Ursgal contributed to an overall lower number of *N*-glycopeptides identified by SugarPy.

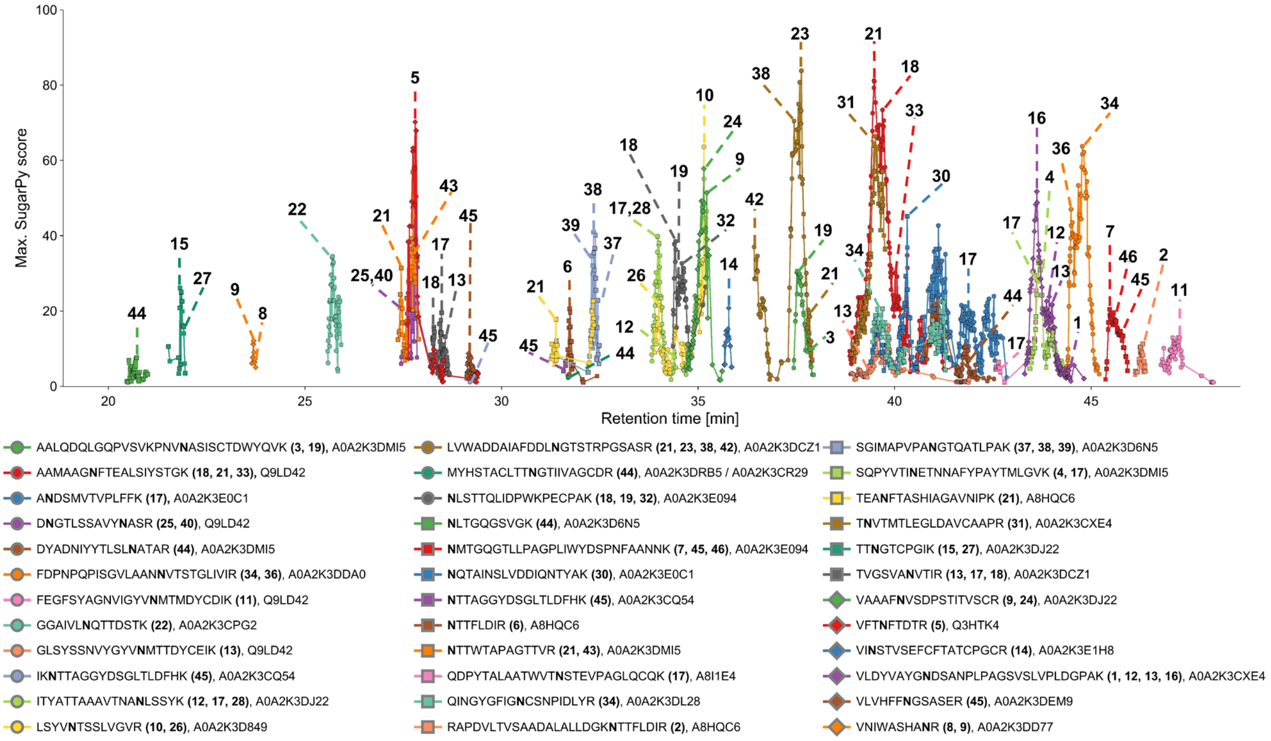


□ N-Acetylhexosamine    Me○ Methylated hexose    ○ Hexose    ☆ Pentose    △ Deoxyhexose

| No. | Composition | Symbols |
|---|---|---|
| 1 | Hex(2)HexNAc(2)MeHex(1)Pent(0)dHex(1) | 0-1x Me○ / 1-5x ○ / 1x △ / 1x ○ / 2x □ |
| 2 | Hex(5)HexNAc(2)MeHex(0)Pent(0)dHex(1) | |
| 3 | Hex(6)HexNAc(2)MeHex(0)Pent(0)dHex(1) | |
| 4 | Hex(3)HexNAc(2)MeHex(1)Pent(1)dHex(0) | 0-1x Me○ / 2-9x ○ / 1x ☆ / 1x ○ / 2x □ |
| 5 | Hex(4)HexNAc(2)MeHex(0)Pent(1)dHex(0) | |
| 6 | Hex(5)HexNAc(2)MeHex(0)Pent(1)dHex(0) | |
| 7 | Hex(6)HexNAc(2)MeHex(0)Pent(1)dHex(0) | |
| 8 | Hex(7)HexNAc(2)MeHex(0)Pent(1)dHex(0) | |
| 9 | Hex(8)HexNAc(2)MeHex(0)Pent(1)dHex(0) | |
| 10 | Hex(10)HexNAc(2)MeHex(1)Pent(1)dHex(0) | |
| 11 | Hex(2)HexNAc(2)MeHex(1)Pent(1)dHex(1) | 0-3x Me○ / 1-6x ○ / 1x ☆ / 1x △ / 1x ○ / 2x □ |
| 12 | Hex(2)HexNAc(2)MeHex(2)Pent(1)dHex(1) | |
| 13 | Hex(3)HexNAc(2)MeHex(1)Pent(1)dHex(1) | |
| 14 | Hex(4)HexNAc(2)MeHex(0)Pent(1)dHex(1) | |
| 15 | Hex(2)HexNAc(2)MeHex(3)Pent(1)dHex(1) | |
| 16 | Hex(3)HexNAc(2)MeHex(2)Pent(1)dHex(1) | |
| 17 | Hex(4)HexNAc(2)MeHex(1)Pent(1)dHex(1) | |
| 18 | Hex(5)HexNAc(2)MeHex(1)Pent(1)dHex(1) | |
| 19 | Hex(6)HexNAc(2)MeHex(0)Pent(1)dHex(1) | |
| 20 | Hex(5)HexNAc(2)MeHex(2)Pent(1)dHex(1) | |
| 21 | Hex(6)HexNAc(2)MeHex(1)Pent(1)dHex(1) | |
| 22 | Hex(6)HexNAc(2)MeHex(2)Pent(1)dHex(1) | |
| 23 | Hex(7)HexNAc(2)MeHex(1)Pent(1)dHex(1) | |
| 24 | Hex(7)HexNAc(2)MeHex(0)Pent(2)dHex(0) | 1x ☆ / 0-1x Me○ / 6-8x ○ / 1x ☆ / 1x ○ / 2x □ |
| 25 | Hex(7)HexNAc(4)MeHex(0)Pent(2)dHex(0) | |
| 26 | Hex(9)HexNAc(2)MeHex(1)Pent(2)dHex(0) | |
| 27 | Hex(1)HexNAc(2)MeHex(3)Pent(2)dHex(1) | 1x ☆ / 0-5x Me○ / 0-8x ○ / 1x ☆ / 1x △ / 1x ○ / 2x □ |
| 28 | Hex(2)HexNAc(2)MeHex(2)Pent(2)dHex(1) | |
| 29 | Hex(1)HexNAc(2)MeHex(4)Pent(2)dHex(1) | |
| 30 | Hex(2)HexNAc(2)MeHex(3)Pent(2)dHex(1) | |
| 31 | Hex(3)HexNAc(2)MeHex(2)Pent(2)dHex(1) | |
| 32 | Hex(4)HexNAc(2)MeHex(1)Pent(2)dHex(1) | |
| 33 | Hex(1)HexNAc(2)MeHex(5)Pent(2)dHex(1) | |
| 34 | Hex(2)HexNAc(2)MeHex(4)Pent(2)dHex(1) | |
| 35 | Hex(2)HexNAc(2)MeHex(4)Pent(2)dHex(1) | |
| 36 | Hex(3)HexNAc(2)MeHex(3)Pent(2)dHex(1) | |
| 37 | Hex(4)HexNAc(2)MeHex(3)Pent(2)dHex(1) | |
| 38 | Hex(5)HexNAc(2)MeHex(2)Pent(2)dHex(1) | |
| 39 | Hex(6)HexNAc(2)MeHex(1)Pent(2)dHex(1) | |
| 40 | Hex(6)HexNAc(4)MeHex(1)Pent(2)dHex(1) | |
| 41 | Hex(7)HexNAc(2)MeHex(0)Pent(2)dHex(1) | |
| 42 | Hex(7)HexNAc(2)MeHex(1)Pent(2)dHex(1) | |
| 43 | Hex(9)HexNAc(2)MeHex(1)Pent(2)dHex(1) | |
| 44 | Hex(4)HexNAc(2)MeHex(0)Pent(0)dHex(0) | 3-5x ○ / 1x ○ / 2x □ |
| 45 | Hex(5)HexNAc(2)MeHex(0)Pent(0)dHex(0) | |
| 46 | Hex(6)HexNAc(2)MeHex(0)Pent(0)dHex(0) | |

**Fig. S3. Identification of tryptic *N*-glycopeptides with MeHex, Pent and dHex modifications from *C. reinhardtii* using the IS-CID/Sugary approach.** A representative elution profile of *N*-glycopeptides identified by SugarPy from IS-CID measurements is shown. Traces represent the maximum SugarPy score of all *N*-glycopeptides for each MS1 spectrum for the corresponding glycopeptide sequence, but glycans are annotated only if they were accepted after manual filtering. *N*-Glycosites identified by protein database search are shown in bold letters. *N*-Glycan compositions reported by SugarPy are given in the table-legend, grouped according to the presence or absence of *N*-glycan decorating Pent and/or dHex. It should be noted that the interpretation of the 146 Da mass increment in corresponding MS1 spectra as dHex is based on previous reports describing *N*-glycosylation in *C. reinhardtii*, while methylated xylose or arabinose, recently identified in other microalgae, might result in the same mass increment (Mathieu-Rivet et al. 2013; Oltmanns et al. 2020; Mocsai et al. 2020). Furthermore, while the IS-CID/SugarPy approach does not reveal glycan structures, the depicted symbol representation of the glycan compositions separates monosaccharides that can be attributed to the glycan core (below horizontal line, based on MS1 fragmentation pattern) from the remaining monosaccharides.

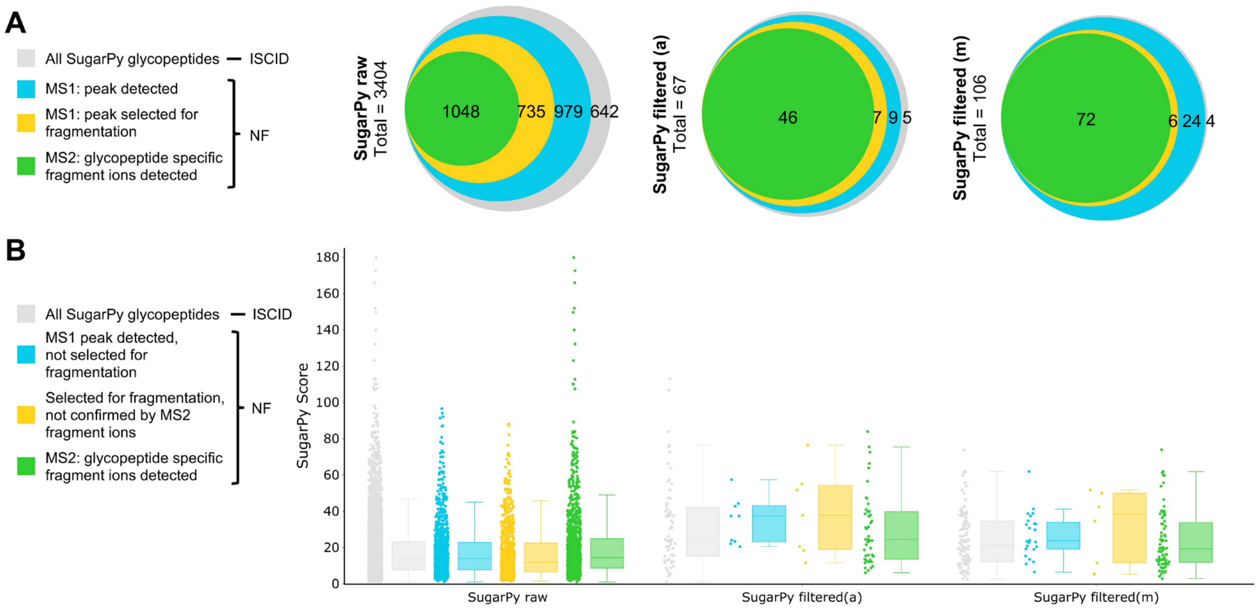


**Fig. S4. Analyses of intact *N*-glycopeptides from *C. reinhardtii* supernatants demonstrate that uncommon *N*-glycopeptides are reliably identified through IS-CID/SugarPy.** (**A**), Initial raw results (top) as well as automatically (middle) and manually (bottom) filtered SugarPy results based on IS-CID measurements (grey) were evaluated through NF measurements. The presence of isotope envelopes on MS1 level corresponding to identified *N*-glycopeptides (blue), their selection for HCD fragmentation (orange) and the detection of glycopeptide-specific fragment ions on MS2 level (green) is shown as area-proportional Venn diagrams. (**B**), The SugarPy score distribution of all identified, unfiltered *N*-glycopeptides (left), automatically (middle), and manually (right) filtered results is represented as boxplots. For each category, the set of all corresponding *N*-glycopeptides (grey), as well as *N*-glycopeptides that were selected for HCD fragmentation and either lacked (orange) or showed (green) the presence of corresponding oxonium and Y-ions on MS2 level, is shown. It should be noted that due to the relatively high number of isomeric combinations (in comparison to human *N*-glycans), the potential for false positives is increased. It can also lead to ambiguous glycopeptide spectrum matches, which are likely to be removed by the employed filtering, even if they showed high SugarPy Scores.

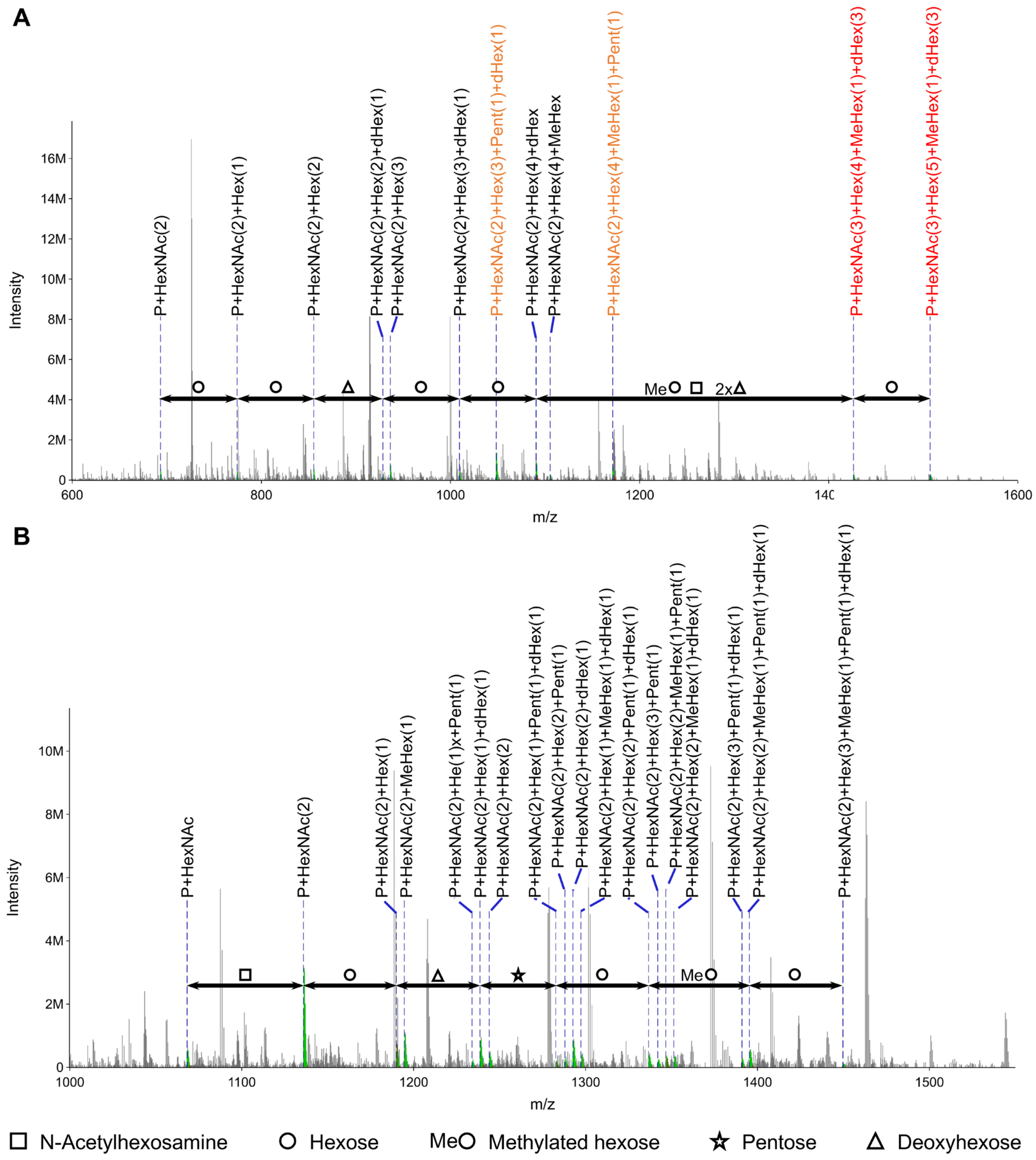


**Fig. S5. Annotated glycopeptide Y-ion series for automatically filtered SugarPy results for *C. reinhardtii.*** The MS1 Y-ion series of glycan compositions Hex(5)HexNAc(3)MeHex(1)dHex(3) (**A**) and Hex(3)HexNAc(2)MeHex(1)Pent(1)dHex(1) (**B**) attached to the tryptic peptides NTTFLDIR (z=2) and VLDYVAYGNDSANPLPAGSVSLVPLDGPAK (z=3), respectively, is shown. These glycopeptides were included in automatically filtered SugarPy results from IS-CID runs, but glycopeptide-specific fragment ions could not be found in NF measurements. Raw peaks for IS-CID-derived spectra are shown in grey, matched peaks in green and annotations as dashed blue lines for the monoisotopic peak. While a continuous Y-ion series could be detected for (**B**), a gap of more than two monosaccharides is apparent in the Y-ion series of (**A**). The Y-ions highlighted in red are therefore not well supported by the fragmentation pattern, while Y-ions marked in orange represent an alternative annotation consistent with the fragmentation pattern.

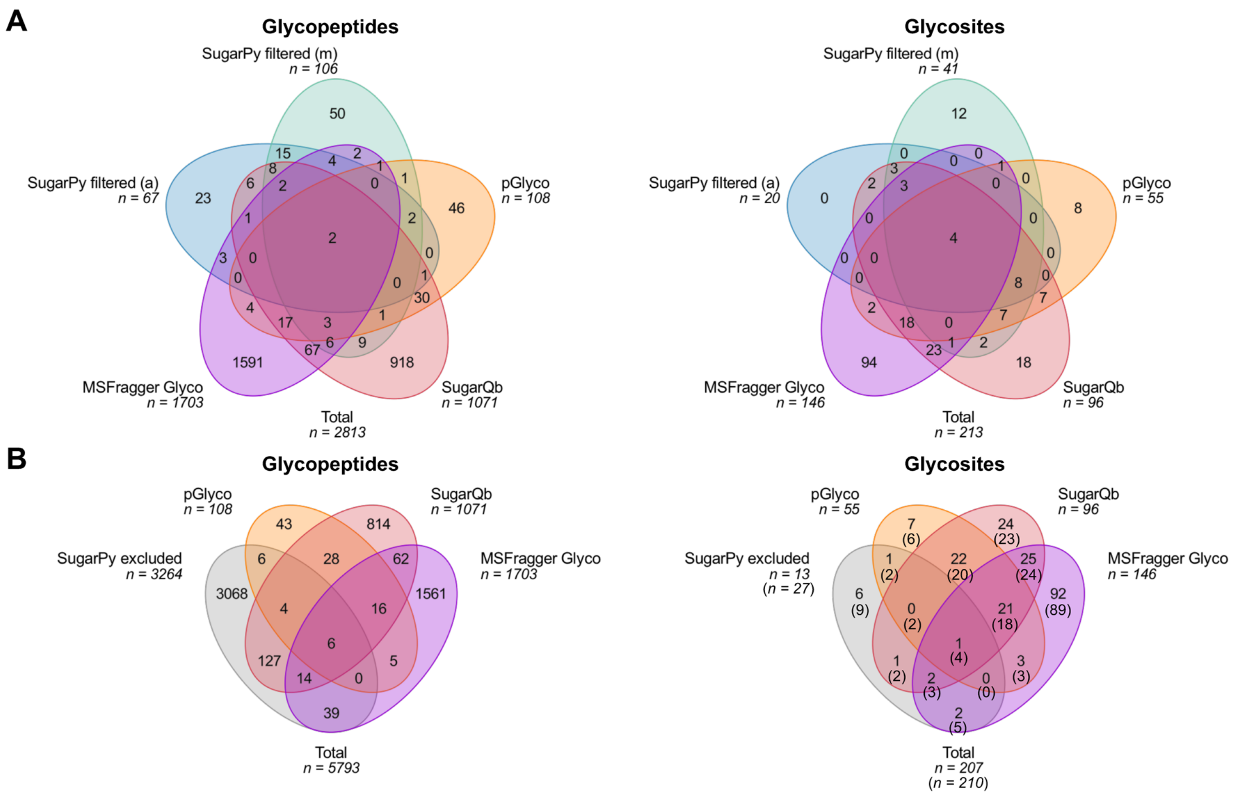


**Fig. S6. *N*-Glycan database dependent algorithms are not suitable for the analysis of *N*-glycopeptides from *C. reinhardtii*. (A),** Venn diagrams for *N*-glycopeptides (defined by the combination of peptide sequence, modification and glycan composition) identified by the IS-CID/SugarPy approach with automatic (blue) and manual (green) filtering or by pGlyco (orange), SugarQb (red), and MSFragger Glyco (purple) from NF measurements, illustrate that only a small *N*-glycopeptide subset (consisting of oligomannosidic *N*-glycopeptides) is found by all three algorithms. Furthermore, large part of unique *N*-glycopeptide identifications from SugarQb and MSFragger Glyco were considered likely false positives, since 39% and 58% of them, respectively, contained ≠2 HexNAc (for one glycosite). (**B**), Comparison of *N*-glycopeptides that were rejected by automatic and manual filtering (grey) with pGlyco and SugarQb results. (**C**) and (**D**), The Venn diagrams of *N*-glycosites for the same categories as in A and B, respectively, shows a higher conformity between the three engines, illustrating that especially for SugarQb and MSFragger Glyco the assignment of *N*-glycan compositions is questionable rather than the identification of *N*-glycosites. Numbers in brackets refer to the identification of putative *N*-glycosites by protein database search with Ursgal, which not always led to the identification of *N*-glycopeptides on MS1 level.

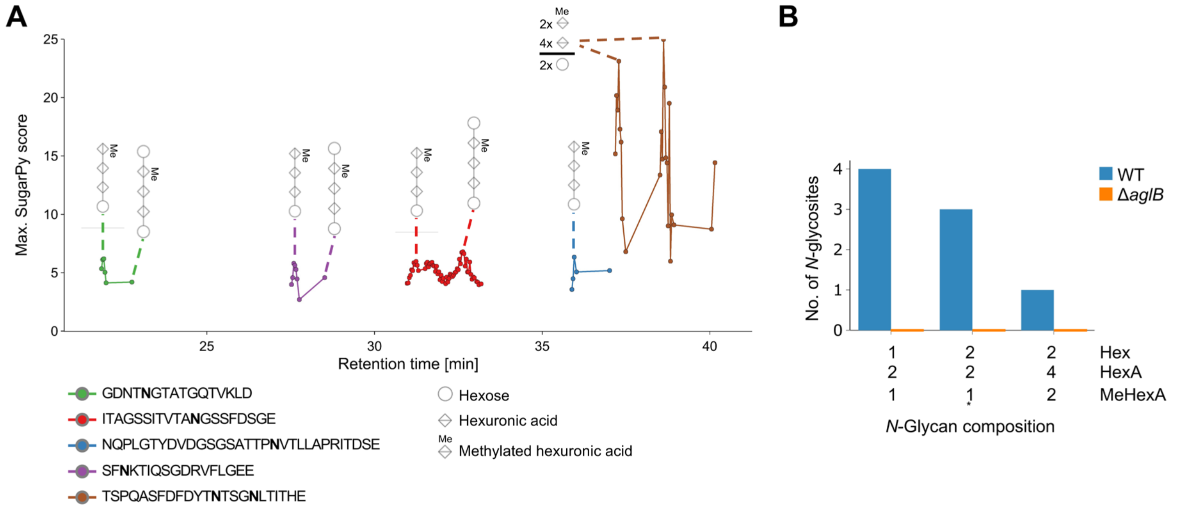


**Fig. S7. SugarPy is suitable for the identification of archaeal *N*-glycopeptides.** (**A**), A representative elution profile of GluC-derived *N*-glycopeptides identified by SugarPy from IS-CID measurements of *H. volcanii* samples is shown (measurements belong to the PRIDE dataset PXD011015). While SugarPy reports glycan compositions, the depicted schematic structural representations (without linkage type) can be deduced from the MS1 fragmentation pattern and are in agreement with the *N*-glycosylation pathway of *H. volcanii*. *N*-Glycosites identified by protein database search are shown in bold letters. For glycopeptides containing multiple glycosites, only combined glycan compositions can be determined. (**B**), All *H. volcanii* *N*-glycopeptides identified with SugarPy after automatic filtering for the wildtype (WT, blue) are supported by literature (*except the glycan composition Hex(1)HexA(3)MeHex(1) which is isobaric with Hex(2)HexA(2)MeHexA(1), resulted in similar SugarPy scores for the same spectra and was therefore removed). The absence of *N*-glycopeptide identifications for samples from the oligosaccharyltransferase knockout mutant Δ*aglB* (orange) supports the reliable functionality of SugarPy.

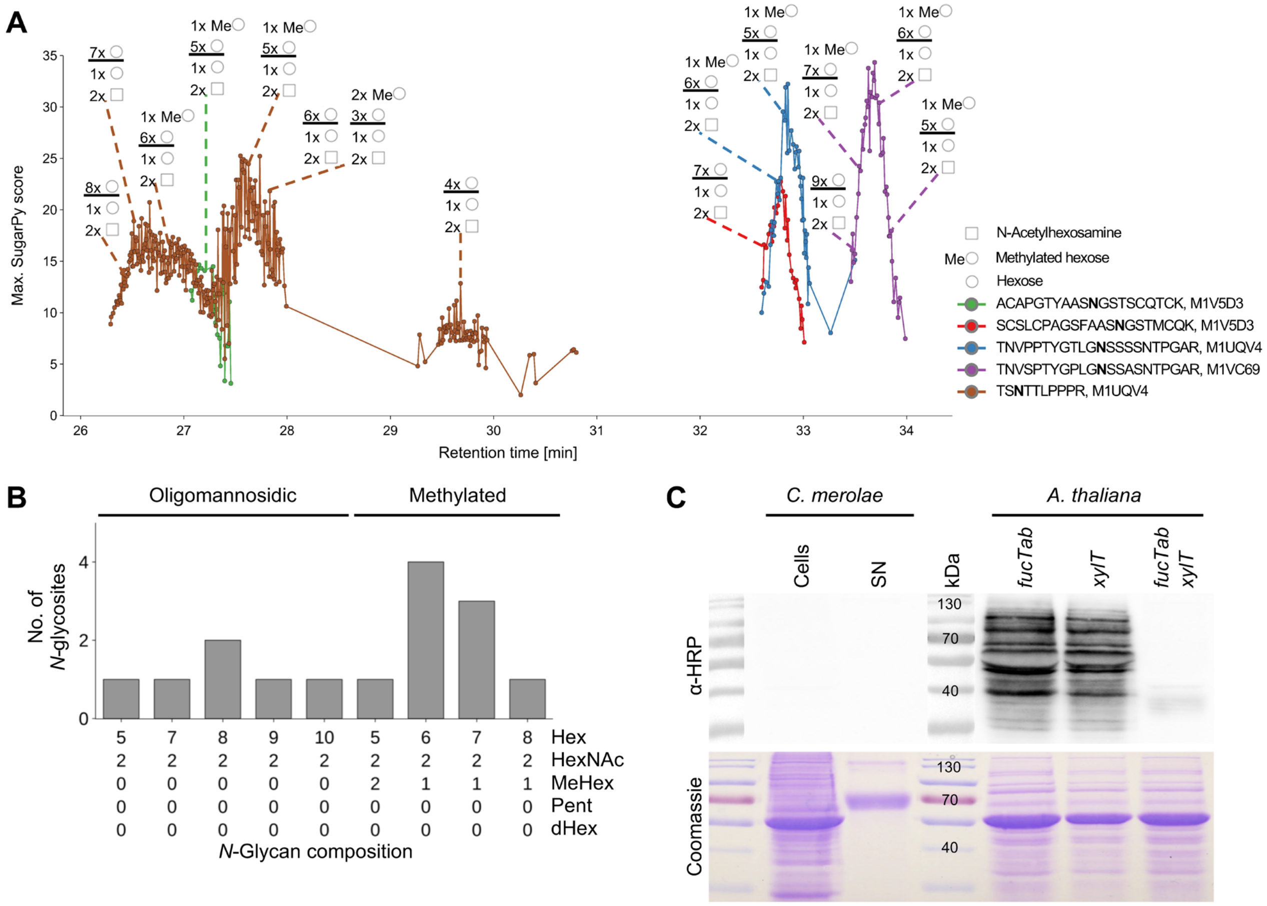


**Fig. S8. *N*-glycans in *C. merolae* are oligomannosidic and partly methylated but devoid of *N*-glycan core modifications.** (A), A representative elution profile of tryptic *N*-glycopeptides identified by SugarPy from IS-CID measurements is shown. For *N*-glycan compositions reported by SugarPy, a common *N*-glycan core is suggested by the observed Y-ions. However, the structure of the *N*-glycan remains to be shown. *N*-glycosites are shown in bold letters and UniProt IDs are given next to the peptide sequence. (B), For each of the *N*-glycans identified from *C. merolae* supernatant samples with SugarPy and manual filtering of results, the number of *N*-glycosites harboring the *N*-glycan is given. Results of three biological replicates are merged. (C), In support of MS analyses, a Western blot using polyclonal anti HRP antibodies (detecting β1,2-xylose and α1,3-fucose residues attached to the *N*-glycan core) showed no signal for total cell extracts or culture supernatant (SN) samples from *C. merolae*, while a strong signal for *A. thaliana* leaf extracts was detected. Sample loading is indicated by a second SDS PAGE stained with Coomassie brilliant blue.

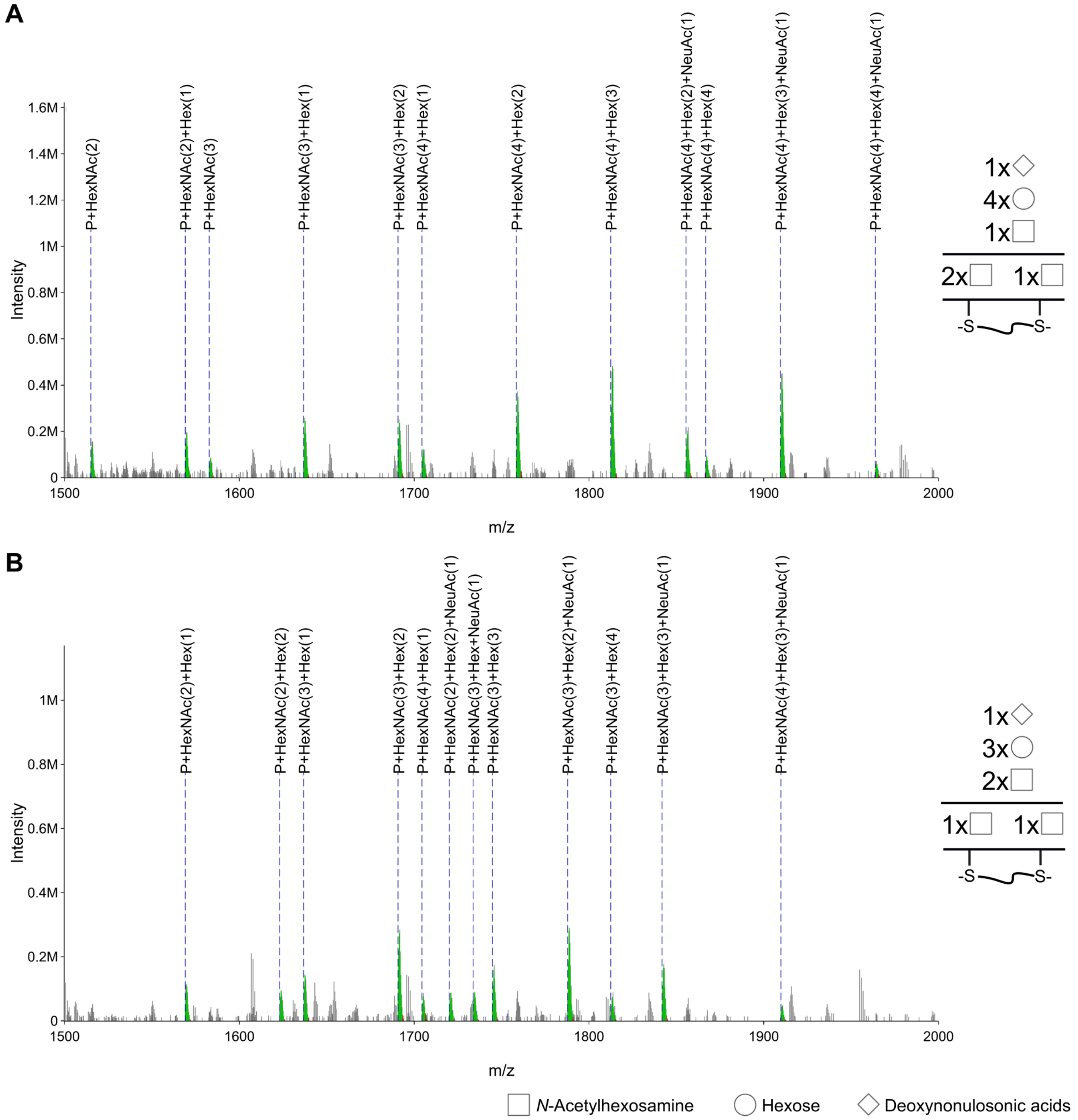


**Fig. S9. IS-CID/SugarPy can be used to identify *O*-glycopeptides.** The MS1 Y-ion series of glycan compositions Hex(4)HexNAc(4)NeuAc(1) (**A**) and Hex(4)HexNAc(3)NeuAc(1) (**B**) attached to the tryptic glycopeptide HYTNPSQDVTVPCPVPSTPPTPSPSTPPTPSPSCCHPR is shown. The peptide sequence was identified to be glycosylated by peptide database search with Ursgal and the absence of an *N*-glycosylation consensus sequence implied an *O*-glycosylation, which is in accordance with the Uniprot prediction of *O*-glycosylation at Ser 105, 111, 113, 119 or 121 for the corresponding protein P01876. MS2 fragmentation spectra suggest the presence of two O-glycans, though their exact position cannot be confidently determined. The depicted symbol representation of the glycan compositions separates monosaccharides that can be attributed to the glycan core (below horizontal line, based on MS1 fragmentation pattern) from the remaining monosaccharides. However, glycan compositions could not be attributed to a specific glycosite. Raw peaks are shown in grey, matched peaks in green and annotations as dashed blue lines for the monoisotopic peak.

**Table S1. All identified *N*-glycosites and several *N*-glycopeptides are supported by literature reports.** All entries are given with their UniProt-ID, corresponding protein name and peptide sequence including the position of the identified *N*-glycosylated Asn within the full protein sequence. Literature reporting the identification of the *N*-glycosite in human milk (bold) or other human fluids/organelles is indicated. For all *N*-glycan compositions identified with SugarPy, supporting literature reporting the same *N*-glycopeptide is provided, if applicable. N-Glycosites that were identified on MS2 level, but for which no glycan could be retrieved from MS1 spectra, are marked with "-". Additionally, it is indicated whether *N*-glycopeptides were identified among automatically (a) and/or manually (m) filtered SugarPy glycopeptides. It should be noted that the following studies used PNGase F treatment and therefore identified *N*-glycosites indirectly through deamidation of Asn: Picariello et al. 2008 (63 *N*-glycosites from milk whey) (Picariello et al. 2008), Cao et al. 2017 (91 *N*-glycosites from mature milk whey) (Cao et al. 2017), Cao et al. 2018 (912 *N*-glycosites from milk fat globule membranes) (Cao et al. 2018), Cao et al. 2019 (96 *N*-glycosites from mature milk whey) (Cao et al. 2019). In contrast, intact *N*-glycopeptides were analyzed by Huang et al. 2017 (111 *N*-glycopeptides from milk whey) (Huang et al. 2017) and Goonatilleke et al. 2019(114 *N*-glycopeptides from milk whey) (Goonatilleke et al. 2019).

| ID | Protein name | Sequence | *N*-Glycosite literature | *N*-Glycan composition | N-glycopeptide literature | SugarPy filter |
|---|---|---|---|---|---|---|
| P01024 | Comp. C3 | TVLTPATNHMG**N**$^{85}$VTFTIPANR | Zhang et al., 2003 | Hex(5)HexNAc(2)NeuAc(0)dHex(0) | - | m |
| | | | **Cao et al., 2019** | | | |
| P01033 | Metalloproteinase inhibitor 1 | FVGTPEV**N**$^{53}$QTTLYQR | Jia et al., 2009 | Hex(4)HexNAc(3)NeuAc(1)dHex(1) | - | m |
| | | | | Hex(5)HexNAc(4)NeuAc(0)dHex(2) | - | m |
| P01591 | IgJ | E**N**$^{71}$ISDPTSPLR | Jia et al., 2009 | Hex(3)HexNAc(3)NeuAc(0)dHex(0) | - | m |
| | | | **Picariello et al., 2008** | Hex(3)HexNAc(2)NeuAc(0)dHex(1) | - | m |
| | | | **Cao et al., 2017** | Hex(5)HexNAc(4)NeuAc(1)dHex(0) | **Huang et al., 2017** | m |
| | | | **Huang et al., 2017** | | | |
| | | | **Goonatilleke et al., 2019** | | | |
| | | | **Cao et al., 2019** | | | |
| P01833 | pIgR (Polymeric immunoglobulin receptor) | QIGLYPVLVIDSSGYVNP**N**$^{186}$YTGR | Ramachandran et al., 2006 | - | - | - |
| | | | **Picariello et al., 2008** | | | |
| | | | **Huang et al., 2017** | | | |
| | | VPG**N**$^{469}$VTAVLGETLK | Jia et al., 2009 | Hex(5)HexNAc(4)NeuAc(0)dHex(3) | - | a, m |
| | | | **Cao et al., 2017** | Hex(5)HexNAc(4)NeuAc(0)dHex(4) | - | m |
| | | | **Huang et al., 2017** | Hex(5)HexNAc(4)NeuAc(1)dHex(2) | - | a, m |
| | | | **Goonatilleke et al., 2019** | Hex(5)HexNAc(4)NeuAc(1)dHex(3) | - | m |
| | | | | Hex(5)HexNAc(4)NeuAc(2)dHex(1) | **Huang et al., 2017** | m |
| | | | | | **Goonatilleke et al., 2019** | |
| | | | | Hex(4)HexNAc(1)NeuAc(1)dHex(5) | - | a |
| | | W**N**$^{499}$NTGCQALPSQDEGPSK | Ramachandran et al., 2006 | Hex(5)HexNAc(4)NeuAc(1)dHex(1) | - | m |
| | | | **Picariello et al., 2008** | Hex(5)HexNAc(4)NeuAc(0)dHex(2) | - | m |
| | | | **Cao et al., 2017** | Hex(4)HexNAc(3)NeuAc(0)dHex(0) | - | a |
| | | | **Huang et al., 2017** | Hex(5)HexNAc(4)NeuAc(0)dHex(1) | - | a |
| | | | **Goonatilleke et al., 2019,** | Hex(5)HexNAc(4)NeuAc(1)dHex(0) | - | a |
| | | | **Cao et al., 2019** | Hex(4)HexNAc(3)NeuAc(0)dHex(1) | - | a |

| | | | | | | |
|---|---|---|---|---|---|---|
| P01857 | IgG1 | EEQY**N**$^{180}$STYR | Jia et al., 2009<br>**Huang et al., 2017** | Hex(5)HexNAc(4)NeuAc(0)dHex(1) | **Huang et al., 2017** | m |
| P01859 | IgG2 | EEQF**N**$^{176}$STFR | Jia et al., 2009<br>**Cao et al., 2017**<br>**Huang et al., 2017**<br>**Goonatilleke et al., 2019** | Hex(5)HexNAc(4)NeuAc(0)dHex(1) | **Huang et al., 2017**<br>**Goonatilleke et al., 2019** | a, m |
| | | | | Hex(4)HexNAc(4)NeuAc(0)dHex(1) | **Huang et al., 2017**<br>**Goonatilleke et al., 2019** | a |
| | | | | Hex(3)HexNAc(3)NeuAc(0)dHex(1) | - | a |
| P01871 | Immunoglobulin heavy constant mu | YK**N**$^{46}$NSDISSTR | Ramsland et al., 2006<br>**Cao et al., 2017**<br>**Huang et al., 2017** | Hex(3)HexNAc(5)NeuAc(0)dHex(1) | - | m |
| P01876 | Immunoglobulin heavy constant alpha 1 | LAGKPTHV**N**$^{340}$VSVVMAEVDGTCY | Jia et al., 2009<br>**Cao et al., 2017**<br>**Huang et al., 2017**<br>**Cao et al., 2018** | Hex(4)HexNAc(4)NeuAc(0)dHex(1) | - | m |
| P02788 | Lactotransferrin | PFL**N**$^{156}$WTGPPEPIEAAVAR | **Picariello et al., 2008**<br>**Cao et al., 2017**<br>**Huang et al., 2017**<br>**Goonatilleke et al., 2019** | Hex(5)HexNAc(4)NeuAc(1)dHex(2) | **Huang et al., 2017**<br>**Goonatilleke et al., 2019** | m |
| | | TAGWNIPMGLLF**N**$^{497}$QTGSCK | **Cao et al., 2017**<br>**Huang et al., 2017** | Hex(3)HexNAc(3)NeuAc(0)dHex(1) | - | a |
| | | | | Hex(4)HexNAc(3)NeuAc(0)dHex(1) | - | a, m |
| | | | | Hex(5)HexNAc(4)NeuAc(0)dHex(2) | - | a, m |
| | | | | Hex(5)HexNAc(4)NeuAc(1)dHex(2) | - | a, m |
| | | | | Hex(4)HexNAc(4)NeuAc(0)dHex(4) | - | m |
| | | | | Hex(5)HexNAc(4)NeuAc(1)dHex(1) | **Huang et al., 017**<br>**Goonatilleke et al., 2019** | a |
| | | SNLCALCIGDEQGE**N**$^{534}$K | **Cao et al., 2018**<br>**Cao et al., 2017** | - | | - |
| | | **N**$^{642}$GSDCPDKFCLFQSETK | **Picariello et al., 2008** | Hex(5)HexNAc(4)NeuAc(1)dHex(2) | - | a, m |
| | | | | Hex(5)HexNAc(4)NeuAc(0)dHex(3) | - | m |
| P08571 | CD14 | **N**$^{151}$VSWATGR | Liu et al., 2005 | Hex(3)HexNAc(2)NeuAc(0)dHex(0) | - | a, m |
| | | | | Hex(4)HexNAc(3)NeuAc(0)dHex(0) | - | a, m |
| | | | | Hex(4)HexNAc(3)NeuAc(0)dHex(1) | - | m |
| P22079 | Lactoperoxidase | IVGYLNEEGVLDQ**N**$^{212}$R | Ramachandran et al., 2006 | Hex(4)HexNAc(3)NeuAc(0)dHex(0) | - | m |
| | | KPSPCEFI**N**$^{358}$TTAR | **Picariello et al., 2008**<br>**Cao et al., 2018**<br>**Cao et al., 2019** | Hex(3)HexNAc(3)NeuAc(0)dHex(0) | - | m |
| P22897 | Macrophage mannose receptor 1 | WECK**N**$^{104}$DTLLGIK | **Cao et al., 2017**<br>**Cao et al., 2019** | Hex(5)HexNAc(4)NeuAc(1)dHex(2) | - | m |
| | | | | Hex(5)HexNAc(4)NeuAc(0)dHex(2) | - | a |
| | | | | Hex(5)HexNAc(4)NeuAc(0)dHex(3) | - | m |
| | | TAHC**N**$^{1205}$ESFYFLCK | **Picariello et al., 2008**<br>**Cao et al., 2018** | Hex(5)HexNAc(4)NeuAc(1)dHex(2) | - | m |

| | | | | | | |
|---|---|---|---|---|---|---|
| P25311 | Zinc-alpha-2-glycoprotein | FGCEIEN**N**$^{128}$R | Ramachandran et al., 2006<br>Jia et al., 2009<br>**Picariello et al., 2008**<br>**Cao et al., 2017**<br>**Cao et al., 2019** | Hex(5)HexNAc(4)NeuAc(1)dHex(1) | - | m |

**Table S2. Detailed LC-MS/MS parameters for TopN, In-Source CID HCD and Top12 analyses with stepped NCE.**

| | In-Source CID HCD (IS-CID) | | | Not fragmented (NF) TopN | NF TopN with stepped NCE | |
|---|---|---|---|---|---|---|
| | human milk sample | *C. reinhardtii* supernatant | *C. merolae* supernatant | | | |
| **Eluent compositions** | Peptide trapping: 0.05% trifluoroacetic acid (TFA) in ultrapure water (A1), 0.05% TFA in 80% acetonitrile (B1)<br>Peptide separation: 0.1% formic acid (FA) in ultrapure water (A2), 0.1% FA in 80% acetonitrile (B2) | | | | | **LC parameters** |
| **Trap Column** | C18 PepMap 100, 300 µM x 5 mm, 5 µm particle size, 100 Å pore size; Thermo Scientific | | | | | |
| **Peptide trapping (eluents A1+B1)** | 2.5% B1 at 5 µL/min for 5 min | | | | | |
| **Flow rate** | 300 nL/min | | | | | |
| **Separation Column** | Acclaim PepMap C18, 75 µm x 50 cm, 2 µm particle size, 100 Å pore size; Thermo Scientific | | | | | |
| **Gradient for peptide separation (eluents A2+B2)** | 2.5% B2 over 5 min,<br>2.5-45% B2 over 40 min,<br>45%-99 % B2 over 5 min<br>99% B2 over 20 min<br>99%-2.5% over 5 min<br>2.5% over 30 min | | | | | |
| **In-source CID** | 90 eV | 80 eV | 70 eV | off | | **MS1 settings** |
| **Use lock masses** | off | | | | | |
| **Ion mode** | positive | | | | | |
| **Resolution at *m/z* 200 (FWHM)** | 70,000 | | | | | |
| **Chromatographic peak width** | 15 s | | | | | |
| **AGC target** | 3e6 | | | | | |
| **Maximum injection time** | 100 ms | | | | | |
| **Scan range** | 600-3000 m/z | | | | | |
| **Mass tags** | On (±203.07937, ±101.53969, ±67.69312, ±50.76984) | | | off | | |
| **TopN** | 12 | | | | | **MS2 settings** |
| **Resolution at *m/z* 200 (FWHM)** | 17,500 | | | | | |
| **Isolation window** | 2 m/z | | | | | |
| **Lower scan range cutoff** | 150 m/z | | | | | |
| **AGC target** | 1e5 | | | | | |
| **Maximum injection time** | 120 ms | | | | | |
| **Normalized collision energy (NCE)** | 30 | | | | stepped; 20, 30, 40 | |
| **Minimum AGC target** | 1.25e3 | | | | | |
| **Intensity threshold** | 1e4 | | | | | |
| **Charge exclusion** | unassigned, >5 | | | | | |
| **Dynamic exclusion** | 15 s | | | | | |

### Supporting Information 1. Details for the collection, processing and analysis of biological samples

#### *C. reinhardtii* and *C. merolae* culture growth conditions

The *C. reinhardtii* wildtype strain cw92 (CC 503) was grown in Tris Acetate Phosphate medium at 25 °C and 20 µE m-2 s-1as liquid culture shaking at 120 rpm. *C. merolae* was grown in shaken (120 rpm) liquid culture at 25°C under 50 µE m-2 s- in 2x Allens medium. Supernatant samples as well as whole cell extracts were obtained as described previously (Schulze et al. 2017).

#### Immunoblot analyses

*C. merolae* samples (30 µg each) and *A. thaliana* leaf extracts (20 µg, obtained by grounding leaves in 10 mM Hepes (pH= 7.4), 150 mM NaCl, 1 mM DTT, followed by removal of cell debris at 14,000 g, 4°C, 15 min) were subjected to immunoblotting employing a HRP antibody (1:20,000 in TBS-T; P7899, Sigma-Aldrich, St. Louis, MO, USA) as described previously (Oltmanns et al. 2020b). In parallel, an SDS PAGE gel, run as a technical replicate, was stained with Coomassie brilliant blue.

#### Isolation of whey proteins from human breast milk

Mature human breast milk samples from a single, healthy human donor, collected four weeks postpartum, or later, were purchased from Biozol Diagnostica Vertrieb GmbH (Eching, Germany). Thawed milk samples were delipidated by centrifugation at 2700 g, 4 °C for 1 h. The resulting fat layer was penetrated with a spatula and the skim milk decanted through a filter paper. Afterwards, the milk was depleted from casein by adding 60 mM CaCl2 and adjusting the pH to 4.6 with hydrochloric acid. Following 30 min incubation at room temperature under agitation, the skim milk was submitted to ultracentrifugation at 189,000 g, 4 °C for 1 h. The casein-depleted whey fraction (supernatant) was decanted through filter paper and stored at -20 °C.

#### Filter Aided Sample Preparation (FASP)

Filter Aided Sample Preparation (FASP) was performed as previously described (Wisniewski et al. 2009; Schulze et al. 2018) loading 60 µg and 100 µg protein, for *C. merolae* and human milk samples, respectively, onto Amicon ultra centrifugal filters (0.5 mL, 30 kDa MWCO, Millipore). Samples were concentrated to 30 µL by centrifugation at RT and 14k rcf. After disulfide bonds were reduced using 200 µL 100 mM DTT in buffer A (100 mM Tris pH=7,6 supplied with 8 M urea) for 30 min at RT, samples were centrifuged at 14k rcf for 15 min (if not indicated otherwise, following centrifugation steps resemble these settings). After two washing steps with buffer A, cysteine residues were masked with carbamidomethyl residues (100 µL of 50 mM iodoacetamide in buffer A; incubation in the dark for 20 min at RT). Three additional washing steps with 100 µL buffer A were followed by five washing steps with 200 µL 50 mM ammoniumbicarbonate (ABC) buffer (centrifugation at 14k rcf for 10 min). Proteins were digested using trypsin (sequencing-grade modified, Promega, Madison, WI, USA; trypsin to protein ratio of 1/50 to 1/100 in 50 mM ABC buffer) for 16 h at 37°C. Peptides were eluted by centrifugation and washing two times with 50 µL $H_2O$ and afterwards were dried in a vacuum centrifuge and stored at -80 °C until the measurement. Digests were performed for three biological replicates from *C. merolae* samples (60 µg) and two technical replicates of human milk protein (100 µg).

**ZIC-HILIC enrichment of *N*-glycopeptides for human samples**

Glycopeptides obtained after FASP digestion were enriched with ZIC-HILIC SPE pipette tips (Protea Biosciences, Morgantown, WA, USA) as described previously (Neue et al. 2011).

**LC and MS parameters**

Peptides were reconstituted in 2 % (v/v) acetonitrile/0.1 %formic acid in ultrapure water and analyzed on an HPLC-MS system consisting of an Ultimate 3000 RSLCnano HPLC (Thermo Scientific, Bremen, Germany), coupled via a nanospray source to a Q Exactive Plus mass spectrometer (Thermo Scientific, Bremen, Germany). A detailed summary of parameters can be found in Table S2. For *N*-glycopeptide analysis, IS-CID was enabled and mass tags were set to select the 12 most abundant precursor ions showing a mass difference of one HexNAc residue for fragmentation. For the validation of SugarPy results, runs without IS-CID were performed.

**Dataset for samples from *Haloferax volcanii***

Samples from *H. volcanii* have been analyzed previously using IS-CID (Esquivel et al. 2016). Briefly, pilins from *H. volcanii* H53 wiltype strain or oligosaccharyltransferase knockout strain ΔaglB were isolated, digested through FASP and measured on a Q Exactive Plus mass spectrometer (Thermo Scientific). Raw files were downloaded from PRIDE (PXD011015) and reanalyzed with Ursgal and SugarPy here.

**Supporting Information 2. Parameters used for peptide database search.**

Raw files obtained from the MS instrument were converted into the HUPO PSI standard format mzML by MSConvert (as part of ProteoWizard 3.0.19046) (Adusumilli and Mallick 2017). All subsequent steps of the analyses were performed within Ursgal (versions 0.6.5 and 0.6.6), a Python framework for bottom-up proteomics tools (Kremer et al. 2016). If not mentioned otherwise, Ursgal's default parameters have been used. For accessing and converting mzML files, pymzML 2.0 (Kosters et al. 2018) was employed. MS Spectra were then matched against a protein database using the search algorithms X!Tandem (version Vengeance) (Craig and Beavis 2004), MS-GF+ (version 2019.04.18) (Kim and Pevzner 2014), and MSFragger (version 20190222) (Kong et al. 2017). For the corresponding datasets, the following databases were combined with common contaminants (https://www.thegpm.org/crap/) and decoys were generated by peptide shuffling for each protein: the *H. sapiens* reference proteome from Uniprot (UP000005640, downloaded 2018/05/09, containing 71617 proteins), the protein sequences of the *C. reinhardtii* v5.5 gene models (Joint Genome Institute) merged with mitochondrial and chloroplast protein sequences from NCBI databases BK000554.2 and NC_001638 (generated on 2015/01/09, containing 18,941 proteins), the *C. merolae* Uniprot database (UP000007014, downloaded on 2019/11/22, containing 5046 proteins), and the most recent annotation of the *H. volcanii* genome (version 06-JUN-2019, doi: 10.5281/zenodo.3565631, consisting of 4186 proteins). For *H. sapiens*, *C. reinhardtii* and *C. merolae*, protein database search was performed with enzyme set to trypsin allowing for two missed cleavages, a precursor mass tolerance of 5 ppm in each direction, a fragment ion mass tolerance of 20 ppm, and the addition of the following modifications: carbamidomethylation of cysteine (fixed), oxidation of methionine (variable), N-terminal acetylation (variable), HexNAc or HexNAc(2) of asparagine (variable, in two independent searches). For *H. volcanii*, the enzyme was gluc and three missed cleavages were allowed, while precursor and fragment ion mass tolerance were set to 10 ppm and modifications were the same but additionally included the following set of variable modifications (as individual searches): Hex, dHex, pentose (Unimod ID: 1425), glucuronyl (Unimod ID: 54), MeHexA ($C_7H_{10}O_6$), HexHexA (Unimod ID: 1427), sulfoquinovose (SQv, $C_6H_{10}O_7S$), Hex(1)HexA(2) ($C_{18}H_{26}O_{17}$), Hex(1)HexA(3) ($C_{24}H_{34}O_{23}$), Hex(1)HexA(2)MeHexA(1)Hex(1) ($C_{31}H_{46}O_{28}$), Hex(1)HexA(2)MeHexA(1) ($C_{25}H_{36}O_{23}$), $SO_3$Hex(1) ($C_6H_{10}O_8S_1$), $SO_3$Hex(1)Hex(1) ($C_{12}H_{20}O_{13}S_1$), $SO_3$Hex(1)Hex(2) ($C_{18}H_{30}O_{18}S_1$), $SO_3$Hex(1)Hex(2)dHex ($C_{24}H_{40}O_{22}S_1$). Search results were statistically post-processed with Percolator (version 3.4.0) (The et al. 2016) or qvality (version 2.02) (Kall et al. 2009) and filtered by posterior error probability (PEP $\leq$ 1 %). Results from all engines and individual searches were merged and for the identification of *N*-glycosites, results were filtered for peptide sequences containing a modified asparagine within the consensus motif N-X-S/T.

**Supporting Information 3. Example for the calculation of the SugarPy score.**

Taking the examples used in Fig. 1, the calculation of the SugarPy score for the glycans HexNAc(2)Hex(4) (glycan1) and HexNAc(1)Pent(1)NeuAc(2) (glycan2) can be explained as follows. For the fragment ion P+HexNAc(2)Hex(4), we assume a normalized intensity of 0.5 and a pyQms mScore of 0.9. Following equation (1), the length of the vector formed by mScore and intensity would be:

$$VL_{\text{glycan1}} = \sqrt{mScore^2 + {i_{norm}}^2} = \sqrt{0.9^2 + 0.5^2} = 1.03$$

In contrast, for the fragment ion P+HexNAc(1)Pent(1)NeuAc(2), assuming the a normalized intensity of 0.2 and a pyQms score of 0.7, equation (1) would result in the following:

$$VL_{\text{glycan2}} = \sqrt{mScore^2 + {i_{norm}}^2} = \sqrt{0.7^2 + 0.2^2} = 0.73$$

This calculation is performed for each matching fragment ion corresponding to one glycan and the results are summed up, e.g. for the glycan HexNAc(2)Hex(4), seven fragment ions were matched.

$$\sum_{i=0}^{n} Vl_{Y_i}(glycan1) = 1.03 + 0.92 + 1.15 + 1.17 + 1.21 + 1.40 + 1.12 = 8$$

For HexNAc(1)Pent(1)NeuAc(2), only three fragment ions were matched.

$$\sum_{i=0}^{n} Vl_{Y_i}(glycan2) = 0.73 + 1.40 + 1.12 = 3.25$$

For the final SugarPy score, the sum of the scores for the individual fragment ion peaks is multiplied by the subtree coverage. The glycan HexNAc(2)Hex(4), fragment ions for seven out of seven potential subtree lengths ($Y_0$ is considered a subtree here) were identified, resulting in a perfect subtree coverage of 1 and a final SugarPy score as follows:

$$STCov_{glycan1} = \frac{N_{subtree_{lengths}}}{L_{total}} = \frac{7}{7} = 1$$

$$SugarPy\ score_{\text{glycan1}} = STCov \times \sum_{i=0}^{n} Vl_{Y_i} = 1 \times 8 = 8$$

In contrast, for HexNAc(1)Pent(1)NeuAc(2), only three fragment ions were matched, resulting in:

$$STCov_{glycan2} = \frac{N_{subtree_{lengths}}}{L_{total}} = \frac{3}{5} = 0.6$$

$$SugarPy\ score_{\text{glycan2}} = STCov \times \sum_{i=0}^{n} Vl_{Y_i} = 0.6 \times 3.25 = 1.95$$

**Supporting Information 4. Parameters employed to run SugarPy.**

The following parameters were set for the sugarpy_run_1_0_0 node within Ursgal for the analysis of all organisms (if not stated otherwise): minimum glycan length: 3, maximum glycan length: 20 (10 for *H. volcanii*), minimum number of spectra: 2, minimum subtree coverage: 0.65 (0.75 for *H. volcanii*), minimum SugarPy score: 1, retention time tolerance: 1 min, precursor mass tolerance plus/minus: 10 ppm, m/z score percentile: 0.7, relative intensity range: 0.2. The mScore cutoff was set to 0.7 for the *H. sapiens* dataset, which analyzed enriched glycopeptides. For all other datasets for non-enriched samples were measured, resulting in lower peak intensities for glycopeptides. This leads to higher relative errors as well as undetectable isotope peaks, both of which result in lower mScores. Therefore, a relaxed mScore cutoff of 0.5 was used to account for lower mScores in these datasets. The following sets of monosaccharides were used for the different datasets: *H. sapiens*: HexNAc, Hex, NeuAc, dHex (resulting in 10625 theoretical glycan molecules); *C. reinhardtii*: HexNAc, Hex, MeHex, dHex, Pent (resulting in 19480 theoretical glycan molecules); *C. merolae*: HexNAc, Hex, MeHex, MeHex(2), NeuAc, dHex, Pent (resulting in 65120 theoretical glycan molecules); *H. volcanii*: HexNAc, Hex, MeHex, NeuAc, dHex, Pent, HexA, MeHexA, dHexN, HexN, SO3Hex, SQv (resulting in 70582 theoretical glycan molecules).

**Supporting Information 5. Detailed procedure for the use of pGlyco, SugarQb and MSFragger Glyco.**

After integration of pGlyco (version 2.2.2) (Liu et al. 2017) into Ursgal, MS raw files were converted using pParse (version 2.2.1 as included in pGlyco) (Yuan et al. 2012) and the glycopeptide search was performed using the glycan database included in pGlyco. Search parameters were the same as described for the protein database search (see above), except no asparagine modification was included and the used protein databases did not contain decoy sequences, since those are generated internally by pGlyco. Furthermore, the FDR calculation of pGlyco was applied and results were filtered by the reported q-value (≤ 1 %).

MSFragger Glyco (version 3.0) (Polasky et al. 2020) was implemented in Ursgal as well, using .mgf input files and the following parameters: msfragger_labile_mode: nglycan; score_ion_list: b, y, b~, y~, Y; intensity_cutoff_diagnostic_ions: 0.1;

min_required_observed_peaks: 15. HexNAc(1), HexNAc(2), HexNAc(2)Hex(1), and HexNAc(1)Hex(1) were included as diagnostic_fragments, while HexNAc(1), HexNAc(2), HexNAc(2)Hex(1), HexNAc(2)Hex(2), and HexNAc(2)Hex(3) were used as modifications_y_ion_offsets.

SugarQb (Stadlmann et al. 2017) was used within Proteome Discoverer 2.1 (Thermo Scientific). The employed workflow consisted of the Spectrum Selector, the MS2 – Spectrum Processor, the G-Score node, SugarQb, Reporter Ion Filtering and the protein database search engines Sequest HT (Eng et al. 1994) and MS Amanda (Dorfer et al. 2014). Parameters were set as listed above and otherwise recommended parameters were used.

The default glycan databases included in pGlyco and SugarQb were used for the analysis of *H. sapiens* samples, while for MSFragger Glyco a human *N*-and *O*-glycan database was downloaded from glySpace using GlycReSoft (Khatri et al. 2016). However, for the analysis of *C. reinhardtii* MS data, a custom glycan database was generated consisting of all theoretical combinations of the monosaccharides HexNAc, Hex, MeHex, Pent and dHex and a maximal glycan length of 20. This theoretical database was used for analyses with SugarQb and MSFragger Glyco. SugarQb and MSFragger Glyco results were filtered by 1% q-value and 1% PEP after post-processing with Percolator and qvality, respectively. Results from all search engines have been sanitized, i.e. for spectra with multiple PSMs, only the best scoring PSM was accepted.

Results from SugarPy, pGlyco, MSFragger Glyco and SugarQb have been compared in regard to identified *N*-glycosites and *N*-glycopeptides. For the latter, monosaccharide modifications of the identified peptide sequence have been removed, since they are included in the glycan composition. Furthermore, glycan compositions reported by the different engines were unified as e.g. HexNac(2)Hex(5)dHex(1), omitting not included monosaccharides (e.g. NeuAc(0)) and the reducing end monosaccharide reported by SugarPy (e.g. End(HexNAc)).
